# Entropic forces drive cellular contact guidance

**DOI:** 10.1101/479071

**Authors:** A.B.C. Buskermolen, H. Suresh, S.S. Shishvan, A. Vigliotti, A. DeSimone, N.A. Kurniawan, C.V.C. Bouten, V.S. Deshpande

## Abstract

Contact guidance—the widely-known phenomenon of cell alignment induced by anisotropic environmental features—is an essential step in the organization of adherent cells, but the mechanisms by which cells achieve this orientational ordering remain unclear. Here we seeded myofibroblasts on substrates micropatterned with stripes of fibronectin and observed that contact guidance emerges at stripe widths much greater than the cell size. To understand the origins of this surprising observation, we combined morphometric analysis of cells and their subcellular components with a novel statistical framework for modelling non-thermal fluctuations of living cells. This modelling framework is shown to predict not only the trends but also the statistical variability of a wide range of biological observables including cell (and nucleus) shapes, sizes and orientations, as well as stress-fibre arrangements within the cells with remarkable fidelity. By comparing observations and theory, we identified two regimes of contact guidance: (i) guidance on stripe widths smaller than the cell size (*w* ≤ 160 μm), which is accompanied by biochemical changes within the cells, including increasing stress-fibre polarisation and cell elongation, and (ii) entropic guidance on larger stripe widths, which is governed by fluctuations in the cell morphology. Overall, our findings suggest an entropy-mediated mechanism for contact guidance associated with the tendency of cells to maximise their morphological entropy through shape fluctuations.

## Introduction

Cellular organization plays a crucial role in the micro-architecture of tissues and dictates their biological and mechanical functioning [1-3]. This organization is often the result of the response of cells to the anisotropy of their micro-environment, which induces cells to migrate preferentially along the direction of anisotropy [4, 5] - a phenomenon called contact guidance [6-8]. Here we make a distinction between directed cell migration that results from contact guidance [4, 5] and the more fundamental contact guidance effect itself, which refers to cell alignment in response to environmental anisotropy [9-11]. Indeed, contact guidance is known to affect various downstream cell behaviours, including survival, motility, and differentiation [11, 12]. Uncovering the origins of cellular contact guidance is therefore critical, not only for understanding tissue morphogenesis and regeneration, but also for predicting disease progression such as cancer invasion [13, 14].

In spite of the importance of cellular organization in all facets of tissue biology, the fundamental question of how a cell organizes itself in physiological tissue context is still poorly understood [15, 16]. Almost exclusively, the rationalisation of contact guidance has been based on the different components of the mechano-transduction pathway. Some of the more common theories hypothesise that: (i) mechanical restriction imposed by the substrate geometry drives the polarisation of linear bundles of F-actin [8], (ii) the tendency of cells to maximise their focal adhesion areas drives their alignment along ridges on grooved substrates [17], and (iii) sharp discontinuities in the substrates (e.g., edges of microgrooves) induce cell alignment by triggering actin polymerisation and thereby focal adhesion formation at these locations [18]. However, these explanations do not provide a general framework for understanding contact guidance (i.e. cannot be extended beyond the particular experimental setup for which they were proposed). A systematic and integrated framework for contact guidance as observed in response to a host of biophysical (microgrooves, substrate elasticity) and/or biochemical (adhesive patterns, ligand density) cues has to-date proved elusive.

The aim of this investigation is to elucidate the general biophysical mechanisms underlying contact guidance of individual cells. Various *in-vitro* chemical micropatterning approaches have been developed to study cellular contact guidance, as model systems to simplify the highly complex *in vivo* environments [19, 20]. We followed such an approach and seeded a low density of myofibroblasts (specifically, Human Vena Saphena Cells; see Methods) on effectively rigid substrates microprinted with fibronectin stripes of widths *w*, ranging from greater than 1.5 mm (resembling a homogeneous 2D substrate) to 50 μm; see Fig. 1a. This enabled us to investigate contact guidance as affected by ECM proteins in the absence of short-range cell-cell interactions [21] and long-range interactions via substrate elasticity [22]. Surprisingly, we observed that even in this highly simplified model system, contact guidance commences at stripe widths much greater than the average cell size. None of the theories mentioned above to rationalize contact guidance, which are based on the interaction of the cell with the edges of the geometrical features used to guide them, can provide an explanation for this observation.

**Figure 1:**
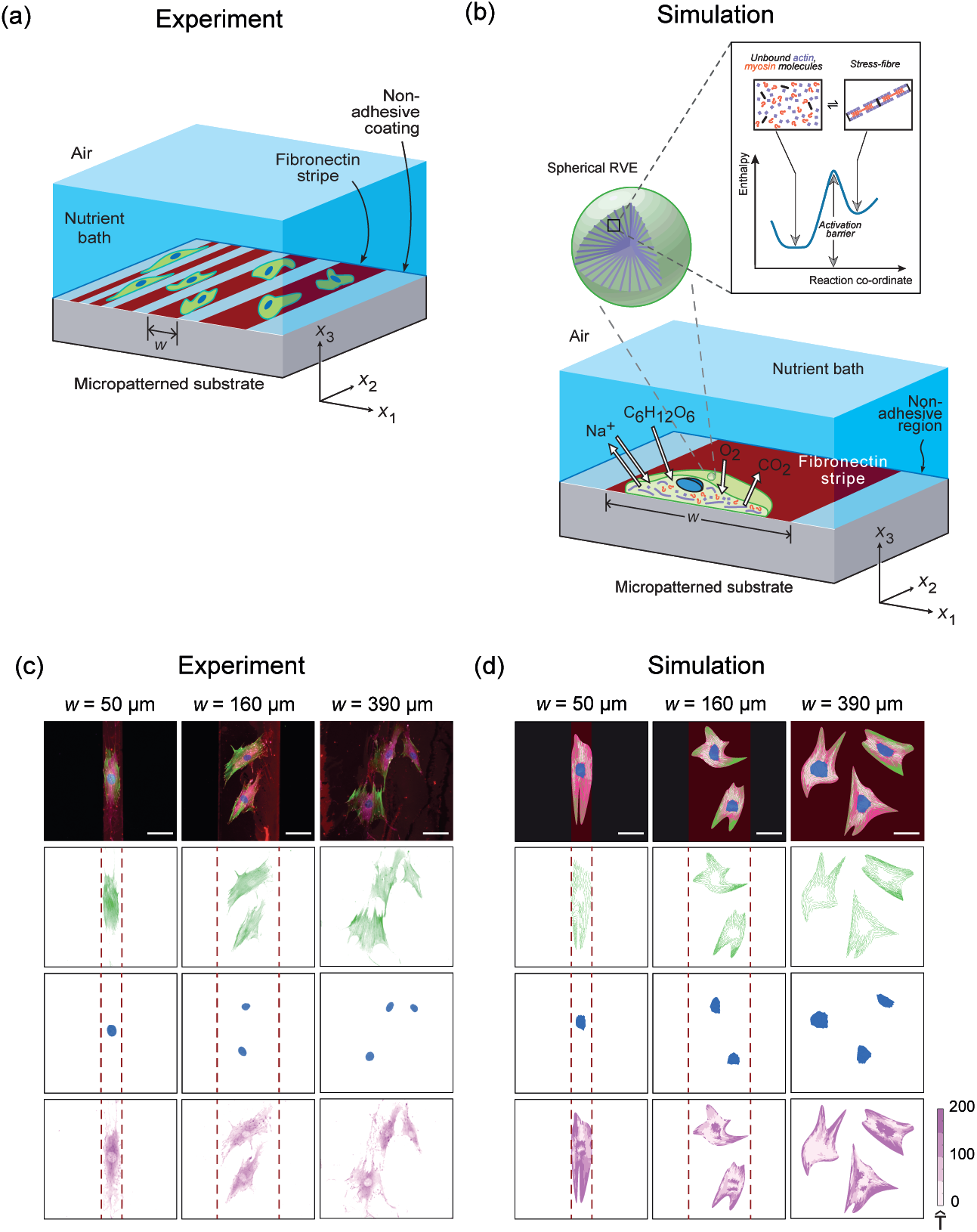
Experimental setup and representative images of the experimental data and simulations. (a) Sketch of the experimental setup of myofibroblasts seeded on a flat substrate micropatterned with fibronectin (maroon) stripes of width *w*. (While each experimental substrate had one stripe width, for conciseness, the sketch shows different stripe widths on the same substrate.), (b) An illustration of the cell model employed in the simulations using homeostatic mechanics framework. The sketch shows a section of a cell on a fibronectin stripe of width *w* and exchanging species with the nutrient bath. The inset shows a representative volume element (RVE) of the cell cytoplasm containing polymerised acto-myosin stress-fibres and the unbound proteins along with the energy landscape that governs the equilibrium of these proteins. (c) Immunofluorescence images of my-ofibroblasts on fibronectin stripes of 50, 160, and 390 µm showing the actin cytoskeleton (green), nucleus (blue), and focal adhesions (magenta); the edges of the stripes are indicated by dashed lines. (d) Corresponding predictions from the homeostatic mechanics framework with focal adhesions parameterised by the magnitude of the normalised traction 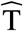 (see Supplementary S2.5.1). The scale bar in (c) and (d) = 60 µm and the width of the fibronectin stripes is indicated by the dashed maroon lines for the two narrowest stripes.

To interpret our unanticipated experimental observations, we combined detailed morphometric analysis of cells (and their subcellular components) with a novel statistical framework that models the non-thermal fluctuations of living cells. These fluctuations, fuelled by the exchange of nutrients between the cell and the surrounding nutrient bath, allow the cell to explore a large number of morphological microstates. However, these fluctuations are constrained by the fact that the cell maintains a homeostatic state and by environmental cues such as, in our case, adherence to fibronectin stripes. In our approach, we assume that the distribution of observed microstates is the one that satisfies these constraints and contains the overwhelming number of morphological microstates (i.e. the distribution that maximises the morphological entropy). Then the statistics of biological observables such as cell shape, orientations and spatial distributions of cytoskeletal proteins is derived from that maximum entropy distribution of morphological states of the cell.

The outcome of our analysis is that, when cells are confined to stripes of finite width, the maximum entropy orientation distribution of cells is not the uniform, but rather one that is peaked around the orientation of the stripe. More specifically, our statistical framework leads to the prediction that there exists a transition stripe width *w*_crit_, above which contact guidance on the stripes is entirely governed by entropic fluctuations of the cell morphology. At stripe widths smaller than *w*_crit_, the confining action of fibronectin stripes causes biochemical changes within the cell that lead to the enhanced polarization of stress-fibres and an increase in contact guidance. In fact, our statistical approach delivers a quantitative measure of the strength of contact guidance through a computable thermodynamic guidance force. This guidance force increases with decreasing *w*, and can be used to identify two distinct regimes of contact guidance, one purely entropic and one that is biochemically-mediated.

A key outcome of this work is that the ordering of cell orientation—a signature of contact guidance—emerges as a consequence of morphological fluctuations of the cell that actually maximise the morphological entropy (or disorder) of cells. Although our results are specific to the phenomenon of contact guidance, our methods are general and they suggest the possibility of a general statistical active-matter-theory for the response of living cells to environmental cues, based on maximisation of morphological entropy subject to the constraint of homeostasis.

## Results

### Cell alignment increases with decreasing stripe width

To investigate the origins of cellular contact guidance, we seeded myofibroblasts on micropatterned fibronectin stripes of width *w*, ranging from 50 µm to homogeneous substrates with *w* → ∞. Outside the stripes, cell adhesion is prevented by treatment with a 1% solution of Pluronic F-127 (see Methods). A schematic illustration of our experimental setup is shown in Fig. 1a, where the fibronectin stripes are represented in maroon and the cells in green. The cells were fixed and stained for actin, vinculin and the nucleus 24 hours after seeding. Representative immunofluorescence images (from at least 50 observations over three independent experiments) of the actin cytoskeleton (green), nucleus (blue), and focal adhesions (vinculin in magenta) of cells on fibronectin (maroon) stripes of width *w* = 50 μm, 160 μm and 390 μm are shown in Fig. 1c (see Fig. S1a for representative observations on all stripe widths investigated here). These images suggest that while the orientation of the actin cytoskeleton and nucleus are reasonably co-ordinated to the cell shape, the focal adhesions are spread nearly evenly throughout the cells. A picture that consistently emerges from these images is that cells align more on the narrower fibronectin stripes than on wider stripes, demonstrating the emergence of contact guidance due to the fibronectin micropatterns.

To further illustrate the effect of substrate micropatterns on cell morphology, randomly selected images of cells are included in Fig. 2a, with the actin stress-fibres shaded in a colour to represent the cell orientation *φ* of the particular configuration. The cell orientation *φ* is defined as the angle between the major axis of the best-fit ellipse to the cell shape and the stripe direction *x*_2_; see Methods. Nearly all cells on the *w* = 50 μm stripes are aligned with the stripes (*φ* ≈ 0°), while there is a much larger spread of cell orientations for cells on wider stripes. Although these images give a flavour of the observations, they do not capture the diversity of the observations, which is best quantified via statistical indicators. The extent of cell alignment to extracellular cues is usually quantified by the cell orientational order parameter Θ (Methods), where Θ = 0 corresponds to a completely random distribution of *φ* and Θ = 1 when all cells have identical orientations [21]. In our experiments, cell alignment steadily increases with decreasing *w*, reaching Θ ≈ 1 for *w* ≤ 160 μm, suggesting complete alignment of the cells with the stripes in this regime (Fig. 2c). Surprisingly, Θ ≈ 0 only occurs in the homogeneous substrate limit (*w* → ∞), while cells align with the stripes for all *w* ≤ 1570 μm that we investigated even though cell sizes (as parametrised by the length of the major axis of the best fit ellipse) are typically on the order of 130 μm.

**Figure 2:**
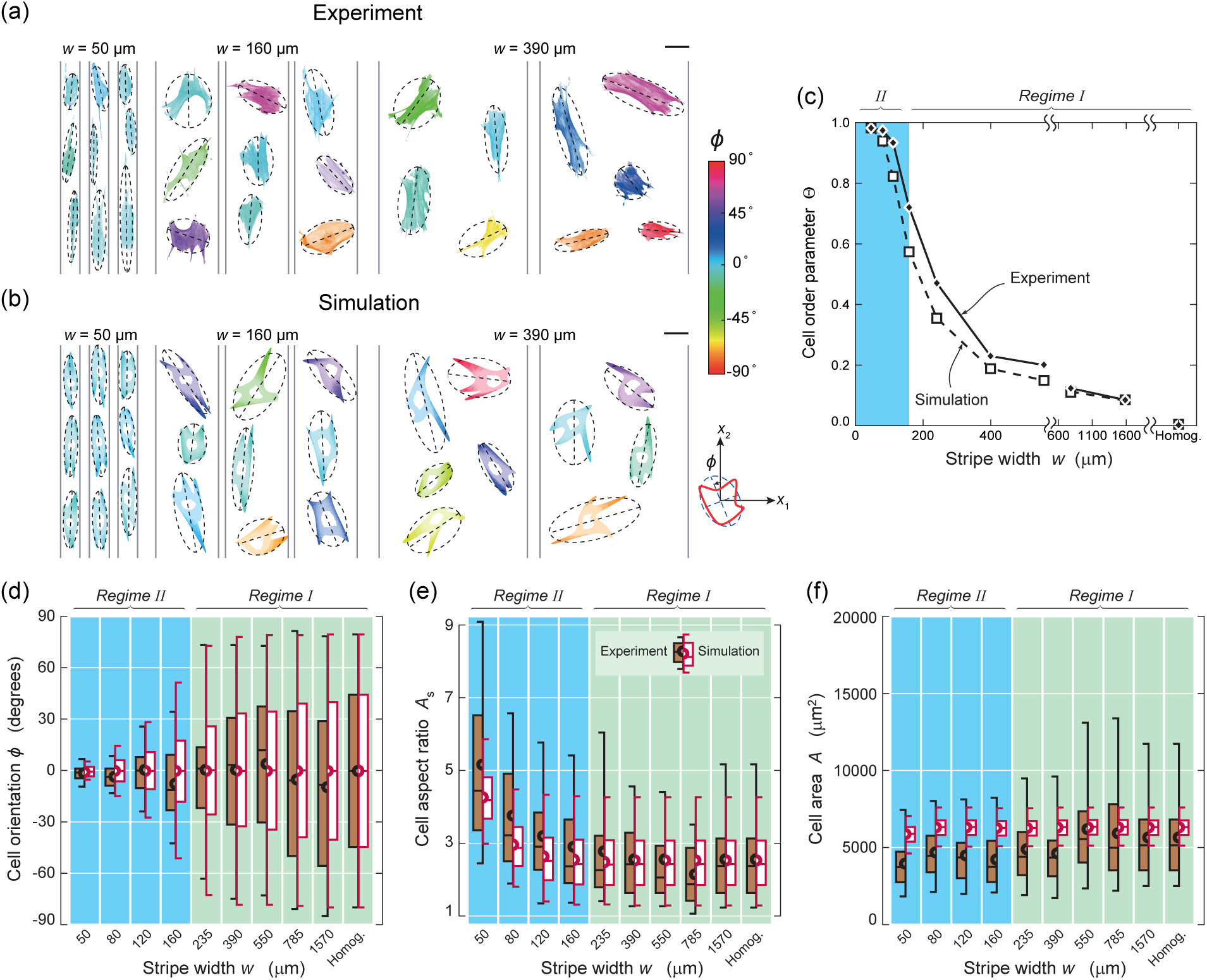
Experimental and computational data on key observables. Images of randomly selected (a) experimentally observed and (b) computationally predicted cell morphologies, showing their actin cytoskeleton and shaded to indicate the cell orientation φ (definition of φ shown in the inset, with the best-fit ellipses and the corresponding major axis indicated on the cell images). The images are shown for cells on three selected widths of fibronectin stripes with the scale bar in (a) and (b) = 60 µm. (c) Measurements and predictions of the cell orientational order parameter *Θ* versus stripe width *w*. Box-and-whisker diagrams of experimental and computational data of the distributions of (d) cell orientation φ, (e) aspect ratio *A*_s_ and (f) area *A* for the range of stripe widths investigated here. The boxes show the quartiles of the distributions with the whiskers indicating the outliers in the experiments and the 5^th^ and 95^th^ percentiles of the distributions in the simulations. The mean of the distributions is depicted by semi-circles for both measurements and simulations. In (c)-(f), the two regimes of contact guidance transitioning at a stripe width *w* 160 µm are indicated and the experimental dataset comprised at least 50 observations per stripe width.

In line with the trend for Θ, the spread in *φ* decreases with decreasing *w* over the entire range of stripe widths investigated here, as shown in the box-and-whisker plots (Fig. 2d). To understand the underlying reason for this spread in cell orientation, we also quantified the effect of the fibronectin stripe width on cell morphology and size. To represent the shape of the cells, we determined the cell aspect ratio *A*_s_ (defined as the ratio between the major axis and minor axis of its best-fit ellipse) and cell area *A* (see Methods). Intriguingly, the dependence of the spread of *φ* on *w* is not reflected in these observables, with the distribution of *A* being independent of *w* whereas the mean *A*_s_ only begins to increase for *w* ≤ 160 μm (i.e., in the regime where Θ ≈ 1); see Figs. 2d-f. These findings underline the need to better understand the origins of the contact guidance response, particularly in a statistical context.

### Homeostatic mechanics predictions reproduce morphometric observations

The experimental system investigated here consists of cells responding to extracellular cues (i.e., adhesive fibronectin stripes), through intracellular processes including cytoskeletal dynamics. The response of this complex system is recorded through a range of observables, all of which exhibit large variations (Fig. 2d-f), but with trends clearly emerging when the statistics of these observables are analysed. This motivates our choice of a statistical modelling framework, which we call homeostatic mechanics, in which, just as in the experimental system, observables fluctuate while trends (and understanding) emerge once these observables are viewed statistically (Supplementary S2). This framework has previously been shown to successfully capture a range of observations for smooth muscle cells seeded on elastic substrates [23], giving us confidence to investigate its generality in terms of predicting the contact guidance behaviour of the myofibroblasts reported here.

The homeostatic mechanics framework recognises that the cell is an open system which exchanges nutrients with the surrounding nutrient bath (Fig. 1b). These high-energy nutrient exchanges fuel large fluctuations (much larger than thermal fluctuations) in cell response associated with various intracellular biochemical processes. However, these biochemical processes attempt to maintain the cell in a homeostatic state, i.e. the cell actively maintains itself out of thermodynamic equilibrium [24] by maintaining its various molecular species at a specific average number over these fluctuations that is independent of the environment [25]. More specifically, homeostasis is the ability of a living cell to maintain, via coupled and inter-connected biomechanical processes, the concentration of all internal species at a fixed average value independent of the environment over all its non-thermal fluctuations. This implies that over the fluctuations of the cell from a fixed reference state (e.g., cell in suspension) 〈Δ*N*_i_〉 = 0 where Δ*N*_i_ is the change in the number of molecules of species *i* from its reference value and 〈x〉 is the average of x over the ensemble of states sampled over the non-thermal fluctuations. These fluctuations alter the cell morphology and each morphological microstate has a unique equilibrium Gibbs free energy *G* = ∑_*i*_ *μ*_*i*_*N*_*i*_, where *μ*_*i*_ is the chemical potential of species *i*. Using the Gibbs-Duhem relation, we then rewrite this in terms of the reference state as 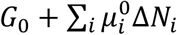 where now 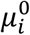 is the chemical potential of species *i* in the reference state. Upon employing the homeostatic constraint that 〈Δ*N*_*i*_〉 = 0, we have 〈*G*〉 = *G*_0_, i.e., irrespective of the environment, the ensemble average Gibbs free energy is equal to that of the cell in suspension. This is a universal constraint that quantifies the fact that living cells maintain themselves away from thermodynamic equilibrium but yet attain a stationary state. Then recognising that biochemical processes such as actin polymerisation and treadmilling provide the mechanisms to explore morphological microstates, we employ the *ansatz* that the observed distribution is that one with the overwhelming number of microstates, i.e. the distribution that maximises the morphological entropy subject to the constraint that that 〈*G*〉 = *G*_0_ and that all microstates are adhered within the fibronectin stripes.

The implementation of the homeostatic mechanics approach described above requires a specific model for the mechano-bio-chemistry of stress-fibres and focal adhesions [26-28]. Here, we employ a relatively simple model, wherein the cell consists of a passive elastic nucleus within a cytoplasm that is modelled as comprising an active stress-fibre cytoskeleton wherein the actin and myosin proteins exist either in unbound or in polymerisation states (Fig. 1b) and elements such as the cell membrane, intermediate filaments and microtubules that are all lumped into a single passive elastic contribution; see Supplementary S2 for details including the cell parameters used to characterise the myofibroblasts.

Figure 1d shows representative images of computed cell morphologies and the corresponding actin, nucleus and focal adhesion organisations (for *w* = 50 μm, 160 μm and 390 μm; directly comparable to observations in Fig. 1c). These morphologies have been selected from the computed ensemble of all morphological microstates the cell samples (i.e. the homeostatic ensemble) wth the constraint that the morphologies have a cell aspect ratio equal to the mode of the distribution for the particular stripe width (see Supplementary S2.5 for the precise definitions of quantities plotted in Fig. 1d and Fig. S1b for corresponding predictions on all stripe widths investigated here). Overall, the cell morphologies are similar to the experimental observations, with the stress-fibres aligning with the cell orientation and the focal adhesions being distributed throughout the cell. Moreover, consistent with the experiments, the cells are randomly oriented on the wider fibronectin stripes but nearly aligned with *φ* ≈ 0° on the *w* = 50 μm stripes (Fig. 2b). To make this comparison more quantitative, we also determine the order parameter Θ from the entire computed homeostatic distribution of cell morphologies, and find excellent agreement with the experimental data over the entire range of *w* (Fig. 2c). Furthermore, the homeostatic mechanics framework accurately reproduces not only the averages, but also the distributions of the observed cell orientation *φ* as quantified in terms of the box-and-whisker diagrams (Fig. 2d). The predictive capability also extended to the cell aspect ratio (Fig. 2e) although now the model under-predicts the spread for the small stripe widths). However, in general the observed spread in the cell areas are larger than model predictions (Fig. 2f). We attribute these discrepancies to the fact that the model assumes a 2D cell (for computational efficiency) and this restricts the spreading of the cell compared to the 3D cell in the experiments. Corresponding morphometric observations and predictions for the nucleus are presented in Supplementary Fig. S2.

### Emergence of two regimes of contact guidance: wide vs. narrow stripes

Both experimental data and model predictions suggest a transition value *w*_crit_ ≈ 160 μm that divides the cell response into two regimes: regime I for *w* > *w*_crit_ where cell alignment occurs in the absence of significant changes of cell shape, and regime II for *w* ≤ *w*_crit_ where cell alignment is accompanied by shape changes (Figs. 2d-f). This transition stripe width is also consistent with the experiments and predictions that cell alignment (as well as nuclear alignment; see Fig. S2) is low for *w* ≫ *w*_crit_ with almost complete alignment (Θ ≈ 1) at widths *w* ≤ 160 μm. Intrigued by this observation, we interrogated the homeostatic mechanics model to evaluate the dependence of total Gibbs free-energy, the cytoskeletal free-energy, and the elastic energy of the cell on stripe width (Fig. S3) as these quantities are typically not directly measurable in experiments. The predictions show that the distributions of cellular energies are invariant to stripe width for *w* > *w*_crit_ and thus hint at the existence of two distinct mechanisms of contact guidance/cell alignment:

i. A regime I, for *w* > *w*_crit_ where cell alignment is accomplished by reorientation of the cell at fixed area and aspect ratio with no associated changes in cellular energies. This suggests that alignment in this regime is not driven energetically but rather by entropic forces (made explicit subsequently) associated with morphological fluctuations of the cell.
ii. A regime II for *w* ≤ *w*_crit_ where cell alignment is accompanied by changes in the cell energies and shapes, and hence thought to be mediated by biochemical changes within the cells.

### Regime I: Entropic alignment of cells for stripe widths larger than the cell size

Cells on substrates with *w* > 160 μm have a mean area and aspect ratio 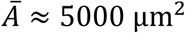 and 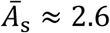 respectively (Fig. 2d). Approximating the shape of the cell to be an ellipse, the semi-major axis *ℓ*_e_ follows 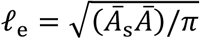. Thus, cells located at the centre of fibronectin stripes of *w* > 2*ℓ*_e_ ≈ 130 μm are expected to assume all orientations *φ* with equal probability, but the orientation of cells on smaller *w* is expected to be restricted by the finite widths of the fibronectin stripes. The critical stripe width of *w*_crit_ = 160 μm is remarkably consistent with 2*ℓ*_e_. Here, we shall use this simple observation to develop an understanding of the mechanisms that result in cell alignment in regime I.

Predictions of the joint probability density function *p*(*x*_c_, *φ*) of the cell centroid being located at position *x*_1_ = *x*_c_ within the stripe and having an orientation *φ* are shown in Fig. 3a-c for three stripes of widths in the range 50 μm ≤ *w* ≤ 390 μm. For *w* = 390 μm, *p*(*x*_c_, *φ*) is nearly uniform, indicating that cells are equally probable to exist at all locations and orientations within the stripe. With decreasing stripe width *p*(*x*_c_, *φ*) starts to become more heterogeneous. Specifically, while cells with centroids near the centre *x*_1_ = 0 of the stripe have equal probability to exist in all orientations, cells near the edges of the stripes are more likely to be aligned with the stripes (*φ* = 0°). This heterogeneity is clearly seen in Fig. 3b where for *w* = 160 μm the probability of misaligned cells (defined arbitrarily here as cells with |*φ*| ≥ 45°) is diminished even for cells with *x*_c_ ≈ 0 and so is the probability of finding a cell near the stripe edges. For the smallest stripe width (*w* = 50 μm), the model predicts that cells are restricted to have their centroids at *x*_c_ ≈ 0 and aligned to the stripes. This transition in the probability distribution occurs at *w* ≈ *w*_crit_ ≈ 2*ℓ*_*e*_ because the major axis of the cell is now longer than the width *w* of the fibronectin stripes and therefore, the cells cannot lie orthogonal to the stripes with |*φ*| = 90°. The model thus suggests that cell alignment in regime I is a direct consequence of the fact that cells near the edges of the stripes necessarily need to be aligned with the stripe directions: with decreasing *w*, the probability of cells being closer to the stripe edges increases, resulting in an increase in the average alignment as parameterised by the order parameter Θ. We refer to this as an entropic alignment, as it is accompanied by no changes to the distribution of the free-energies the cell attains as it fluctuates over its homeostatic state (Fig. S3). Rather, it is only associated with the spatial restriction that the fibronectin stripes impose on the states (or morphologies) the cell is allowed to attain and is therefore entropic in origin.

**Figure 3:**
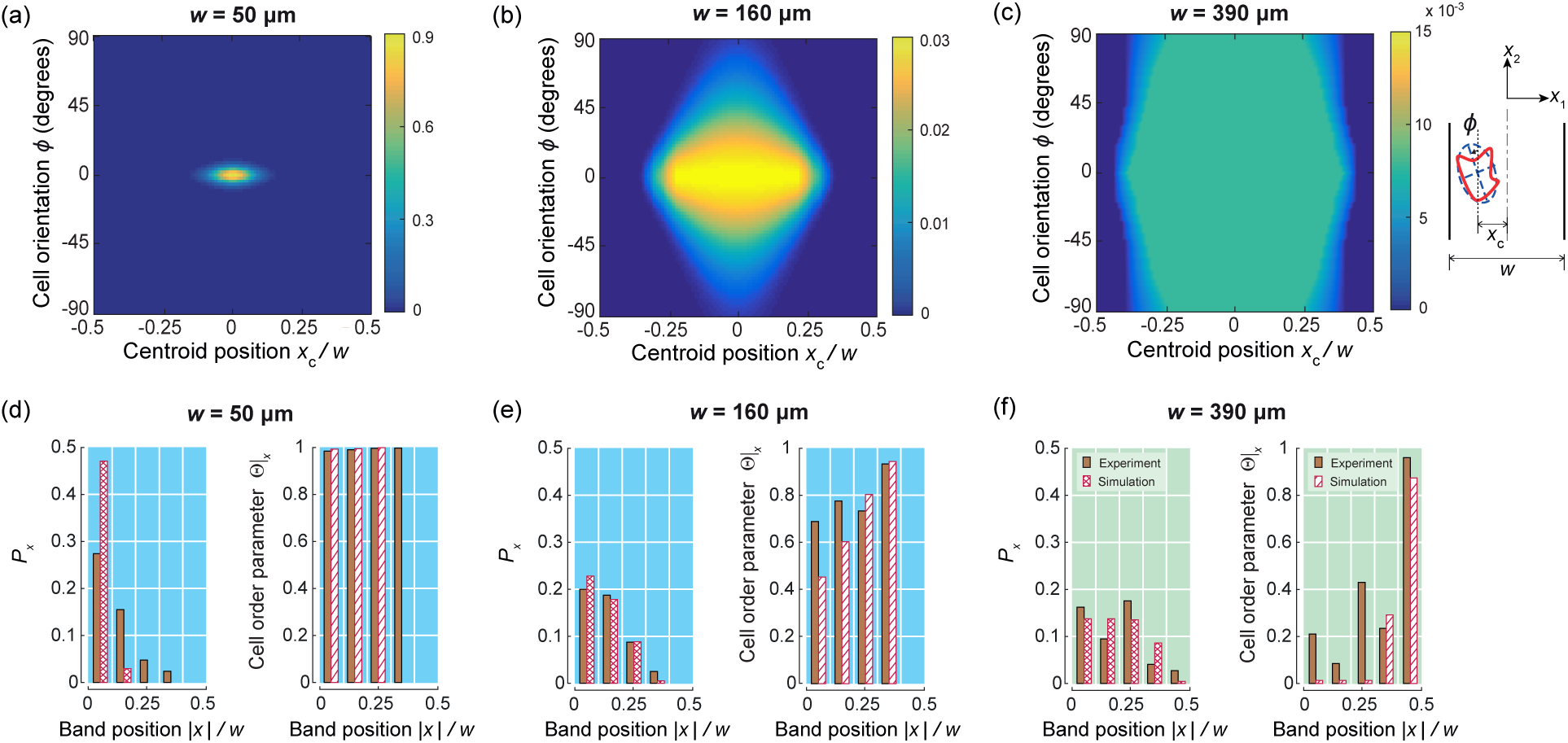
(a-c) Predictions of the joint probability density distributions *p*(*x*_c_/*w*, φ) of the cell centroid being located at position *x*_1_ *= x*_c_ within the stripe and having an orientation φ. Results are shown for three selected stripe widths *w* with the inset defining *x*_c_ and φ. With decreasing *w*, the distributions become more heterogeneous with cells unable to adopt an isotropic distribution of orientations especially near the stripe edges. Note the di*ff*erences in the colour scales of the probability densities between the di*ff*erent sub-parts. (d-f) Measurements (from 50 observations per stripe width) and predictions of the probability *P_x_* of the cell centroid location across the stripe (left) and the conditional order parameter *Θ _x_* of cells within each band centred at location *x* (right). In plots (d-f) each stripe is divided into bands of equal width and the symmetric results shown for |*x*| varying from the stripe centre (*x* = 0) to the stripe edge at |*x*|/*w* = 0.5.

A key assumption in the model is that on the experimental timescales (the 24 hours the cells were cultured on the micropatterned substrates) over which the orientations and positions of the cells fluctuate, the behaviour is “ergodic” (i.e., the cells sample all states compatible with the constraints with equi-probability) with the consequence that the distribution of states they assume maximises their morphological entropy. A corollary of this assumption is that over these timescales the cells have no memory such that the cells might align while near the stripe edges but they lose the memory of that state as they migrate to the interior of the stripes. Based on the entropic *ansatz* that all allowable states are equally probable the model predicts that: (i) cells near the centre of stripes of width *w* > 2*ℓ*_*e*_ are equally likely to exist in any orientation, and (ii) given that cells near the stripe edges exist in fewer orientations, cells are less likely to be found near the stripe edges. To confirm that the experimental results agree with these predictions, we divide each stripe into bands of breadth *w*/10 and measure: (i) the probability *p*_*x*_ of finding a cell in a band with midpoint located at *x*_1_ = |*x*|, and (ii) the corresponding order parameter Θ|_*x*_ of cells located within that band. These measurements for cells on stripes of widths *w* = 50 μm, 160 μm and 390 μm, together with the corresponding model predictions, are included in Figs. 3d-f. Indeed, in both the measurements and predictions the probability of finding a cell in a band near the stripe edge, i.e. |*x*|/*w* ≈ 0.5 is lower than finding the cell at the centre of the stripes. By contrast, while the order parameter Θ|_*x*_ ≈ 1 in all bands for the *w* = 50 μm stripe, the order parameter is higher in bands near the stripe edges compared to bands near the stripe centre for the two larger stripe widths. The consistency and agreement between measurements and predictions strongly suggests the validity of the no-memory assumption, with cell alignment (Fig. 2b) only resulting from those cells that directly interact with the stripe edges.

### Regime II: Biochemical changes accompany alignment at small stripe widths

The enhanced cell alignment in regime II (*w* ≤ 160 μm) is accompanied by changes to cell shape as parameterised by the cell aspect ratio (Fig. 2e). We hypothesise that these shape changes go hand-in-hand with biochemical changes within the cells. To test this idea, we measured the dispersion of actin stress-fibre orientations *φ* within cells and used this distribution as a direct observable indicative of the biochemical state of the cell. Specifically, we monitored local stress-fibre orientations *φ*(*x*_*i*_^)^ with respect to the *x*_2_-direction at every location *x*_*i*_ within the cell and defined a stress-fibre orientation measure 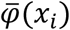 that is invariant to rigid body rotations of the cell (Supplementary S1.2). Figure 4a shows representative images of observed cells with the stress-fibres coloured by their rotationally invariant orientation 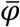 Cells on *w* = 50 μm stripes have a near uniform distribution of stress-fibre orientations with 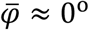 throughout the cell, however, there is a larger dispersion in 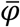 for cells on wider fibronectin stripes. To quantify this observation more precisely, we evaluated probability density functions 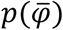 by assembling the measurements of 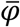 over all imaged cells (at least 50 cells) for each stripe width. The distributions 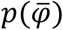 shown in Fig. 4c hardly change in regime I (*w* > 160 μm), but become increasingly peaked with mode at 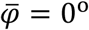 in regime II (*w* ≤ 160 μm).

**Figure 4:**
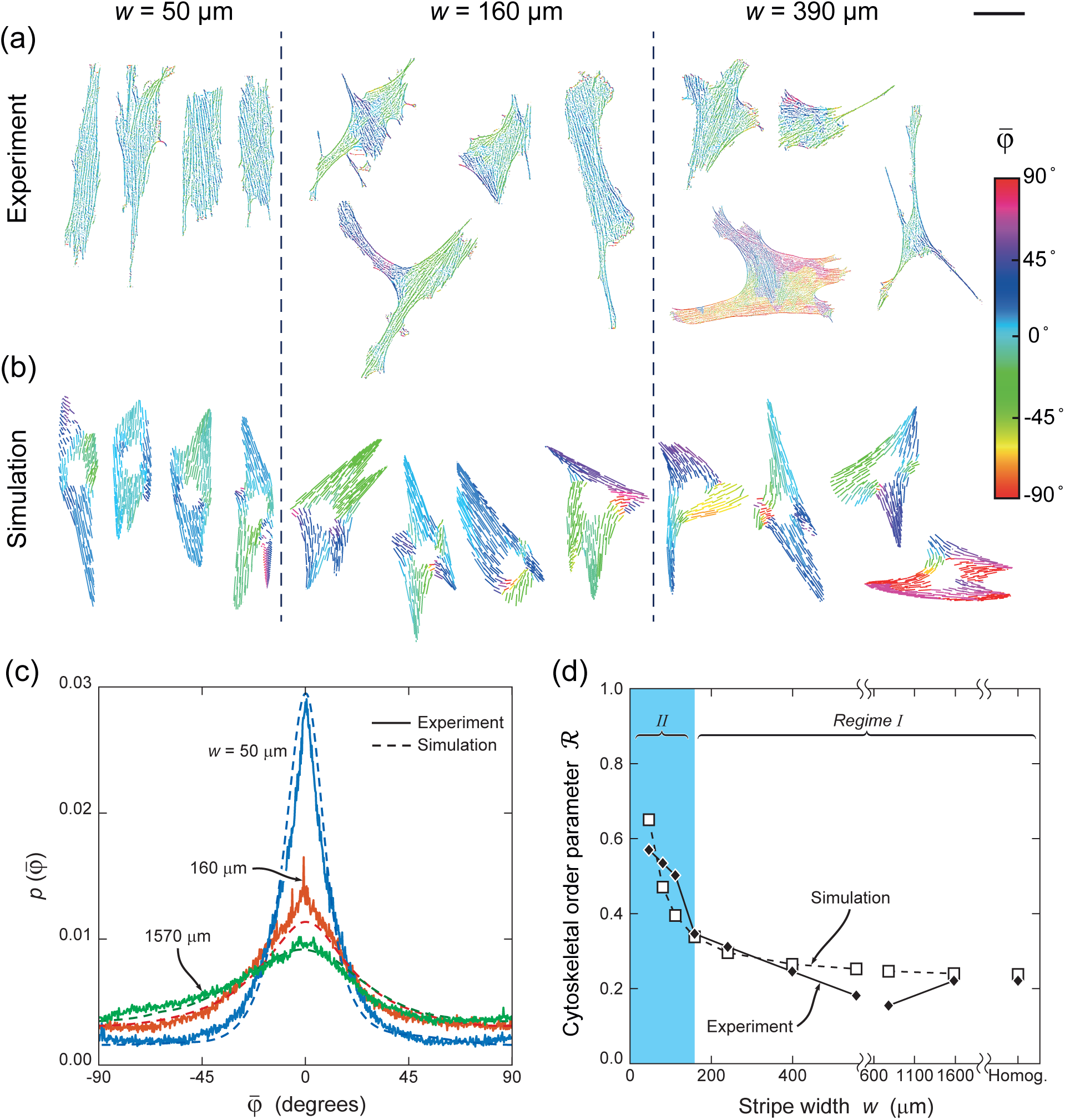
Experimental and computational data for the stress-fibre distributions. (a) Experiments and (b) simulations of the stress-fibre distributions within cells on three stripe widths *w*. The region with no stress-fibres is the passive nucleus in the 2D model. The stress-fibres are coloured by their orientation as parameterised by the measure 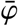 that is invariant to rigid body rotations of the cell. The scale bar in (a) and (b) = 60 µm. (c) The corresponding predictions and measurements of the probability density functions 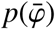 over the ensemble of cell morphologies for three selected stripe widths. (d) Predictions and measurements of the cytoskeletal order parameter *R* extracted from 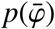 as a function of *w*. The experimental dataset in (c) and (d) comprised 50 observations with both the experiments and simulations illustrating that cytoskeletal changes, as parameterised via stress-fibre ordering *R*, only commence in regime II.

To succinctly capture this decrease in the dispersion of 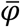, we introduce a cytoskeletal order parameter ℛ (analogous to the cell orientation order parameter Θ) as

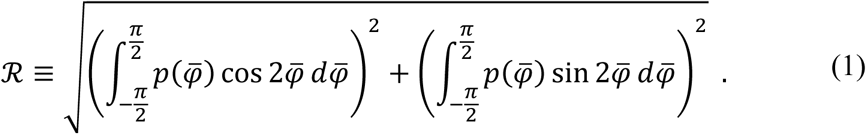

We emphasize that ℛ is a statistic defined over the entire ensemble of observed cells through the distribution 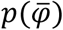 and differs from definitions used in previous studies [29, 30]. This definition not only gives more robust results, but is also more in line with the fluctuating response of cells as proposed by the homeostatic statistical mechanics framework; see detailed discussion in Supplementary S2.5.3-S2.5.4. We find that ℛ, unlike Θ, is approximately constant for stripe widths *w* > 160 μm, but rises sharply for *w* ≤ 160 μm (Fig. 4d). This suggests that no changes to the biochemical state of the cells occur in regime I, but cell alignment is accompanied by biochemical changes in regime II. The corresponding predictions of the spatial distributions of stress-fibre orientations 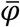, the probability density functions 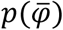, and ℛ are all in remarkable correspondence with detailed experimental measurements of the state of the cells; see Figs. 4b, 4c and 4d, respectively and Supplementary S2.5.3 for details of the calculation of 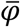 from the model. In particular, both experimental observations and model predictions show negligible enhancement in stress-fibre alignment in regime I, but increasing stress-fibre polarisation with decreasing stripe width in regime II.

### Thermodynamic forces help delineate the two regimes of alignment

Given the high level of agreement between the detailed experimental data and the predictions of the homeostatic framework, we shall now interrogate the model to better understand these regimes. In the model, the coarse-graining of the biochemical processes results in the production of morphological entropy that is conjugated to a scalar parameter 1/*ζ*. By analogy with statistical thermodynamics, we call 1/*ζ* homeostatic temperature. Thus, while ℛ reflects the biochemical state of the cell as indicated by the experimentally-measurable stress-fibre cytoskeleton, 1/*ζ* is expected to capture the *overall biochemical state* of the cells. To illustrate this, we include predictions (Fig. 5a) of the normalised homeostatic temperature 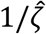(see Supplementary S2.5 for detailed definitions) as a function of stripe width. The homeostatic temperature 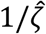 is invariant in regime I, but rises sharply in regime II as cells undergo biochemical changes to maintain a homeostatic state when spatially constrained by narrow fibronectin stripes. Thus, similar to ℛ, the homeostatic temperature shows a clear delineation between the two regimes and we shall use this understanding to employ other quantities suggested by the analogy with thermodynamics to more precisely define the regimes of contact guidance.

**Figure 5:**
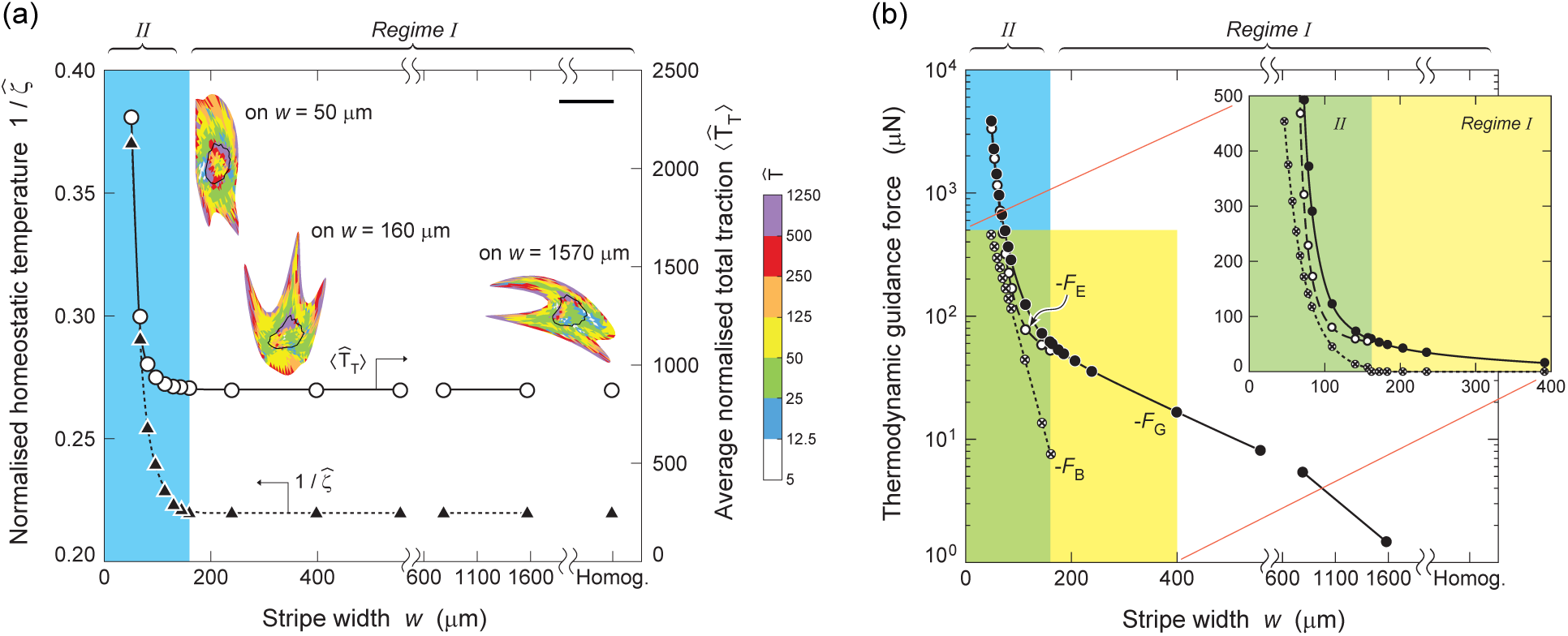
Homeostatic temperature, traction forces and guidance forces. (a) Predictions of the variation of the normalised homeostatic temperature 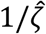 and average normalised total traction force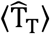 with stripe width *w*. Predictions of the spatial distributions of normalised tractions 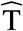 are also included for randomly selected cell morphologies on three stripe widths (outline of nucleus shown as a black line) with the scale bar = 60 µm. Both homeostatic temperature and total traction forces only change with stripe width in regime II. (b) Predictions of the total guidance force *F*_G_ as well as its entropic and biochemical components *F*_E_ and *F*_B_, respectively as a function of *w*. The inset shows the variation using a linear scale for the forces and illustrates that *F*_B_ = 0 for *w* > 160 µm (regime I).

In classical thermodynamics, forces conjugated to the (internal) energy of a system are routinely defined as the force driving the system to reduce its energy [31-33]. Similarly, entropic forces that originate from thermal fluctuations drive the system towards a state that maximises its entropy and are thus conjugated to the entropy of the system [31-33]. These force definitions unambiguously discriminate regimes where the behaviour of a system is governed either energetically or entropically. We shall use a similar strategy to more precisely characterise the two regimes of contact guidance. Recall that equilibrium of a system in contact with a thermal bath at temperature *T* is defined through the canonical ensemble with the thermodynamic driving force *F* given by *F* ≡ (∂*F*/∂*X*)_*T*_. Here, *F* is the Helmholtz potential while *X* is the thermodynamic displacement conjugated to *F*. In an analogous manner, homeostatic statistical mechanics defines equilibrium of a living cell exchanging nutrients with its environment via the homeostatic ensemble, with the homeostatic potential ℳ playing a role similar to the Helmholtz potential. The thermodynamic guidance force constraining the cells to lie within the fibronectin stripes is then given by (see Supplementary S3 for a more detailed definition and corresponding derivations)

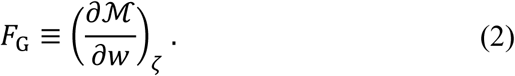

Here, 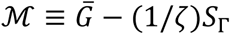, with 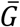 the average Gibbs free-energy of the cells in the homeostatic ensemble at homeostatic temperature 1/*ζ*, while *S*_Γ_ is the equilibrium morphological entropy of the cells. Thus, similar to classical thermodynamic forces, *F*_G_ comprises two components: (i) a biochemical contribution 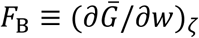 resulting from the propensity of the cell to minimise the average Gibbs free-energy of the morphological states it samples, and (ii) an entropic contribution *F*_E_ ≡ −(1/*ζ*)(∂*S*_Γ_/∂*w*)_*ζ*_ from the tendency of the cell to maximise its morphological entropy. Predictions of these forces as a function of stripe width (Fig. 5b) illustrate that *F*_E_ is the dominant contributor to *F*_G_ over the full range of stripe widths, although *F*_B_ ≈ *F*_E_ for the narrowest stripes (*w* = 50 μm). Importantly, *F*_B_ = 0 for *w* > 160 μm (inset of Fig. 5b), implying that only entropic forces contribute to the alignment of cells within these fibronectin stripes. This unambiguously defines regime I as a purely entropic regime. Also, since both *F*_E_ and *F*_B_ ≠ 0 in regime II (*w* ≤ 160 μm), with their magnitudes being similar for the narrowest stripes, we refer to regime II as biochemically-mediated.

### Orientational alignment is a consequence of the tendency of cells to maximise morphological disorder

In regime I, the leading cell dimensions are smaller than the width of the fibronectin stripes. Cells in this regime thus have the possibility to be completely orientationally disordered (Θ = 0), but “choose” to orientationally order and align with the stripes. This alignment occurs because cells adhered near the stripe edges necessarily need to be aligned with the stripe direction. The question then remains as to why cells wander towards the stripe edges which in turn leads to an apparent decrease in their entropy (disorder). This behaviour of individual cells is reminiscent of orientational ordering in a fluid of hard rods. In his study of a fluid of hard rods, Onsager [34] showed that while the orientational entropy of the ordered phase is lower than that of the isotropic phase, the total entropy of the orientationally ordered phase is higher than the isotropic (orientationally disordered) phase. This is because the orientationally ordered phase has a higher translational and hence total entropy due to the decrease of excluded volume caused by alignment. In other words, the system has traded-off translational and orientational entropies so as to maximise its total entropy. We can extrapolate this idea to understand how the non-thermal fluctuations of individual cells results in them being orientationally ordered when seeded on the fibronectin stripes.

To illustrate, in a simplistic manner, how this mechanism could be at play in our system, we examine a hard rod of length *L* in a channel^2^ of width *w*, with no part of the rod allowed to penetrate the channel edges (Supplementary S4). Three models are analysed: (i) the full model (FM) where the rod has both translational and rotational freedom; (ii) the translationally constrained (TC) model where the rod centroid is pinned at the channel centre *x*_1_ = 0; and (iii) the orientationally constrained (OC) model where the rod is constrained to be aligned with the channel (*φ* = 0). Illustrations of the states the rod assumes in three channel sizes 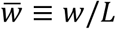 are included in Fig. 6a for the full model. For the smallest channel size, the orientations of the rod are highly constrained, since the rod is not able to explore an isotropic distribution of orientations (−π/2 ≤ *φ* ≤ π/2). By contrast, for 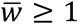, the rod assumes an isotropic distribution of orientations when located near the centre of the channel with the fraction of the channel width over which such isotropic distributions are possible increasing as 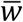 becomes larger. As a consequence, the orientational order parameter Θ_R_ of the rod (analogous to the orientational order parameter Θ of the cell) increases with decreasing 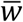(Fig. 6b), in a manner similar to that for cells on fibronectin stripes (Fig. 2c). By contrast, when the rod is translationally constrained, ordering occurs only for 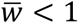 as the rod is unable to wander to the edges of the channel where it necessarily needs to align. Finally, when the rods are orientationally constrained, Θ = 1 for all 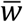.

**Figure 6:**
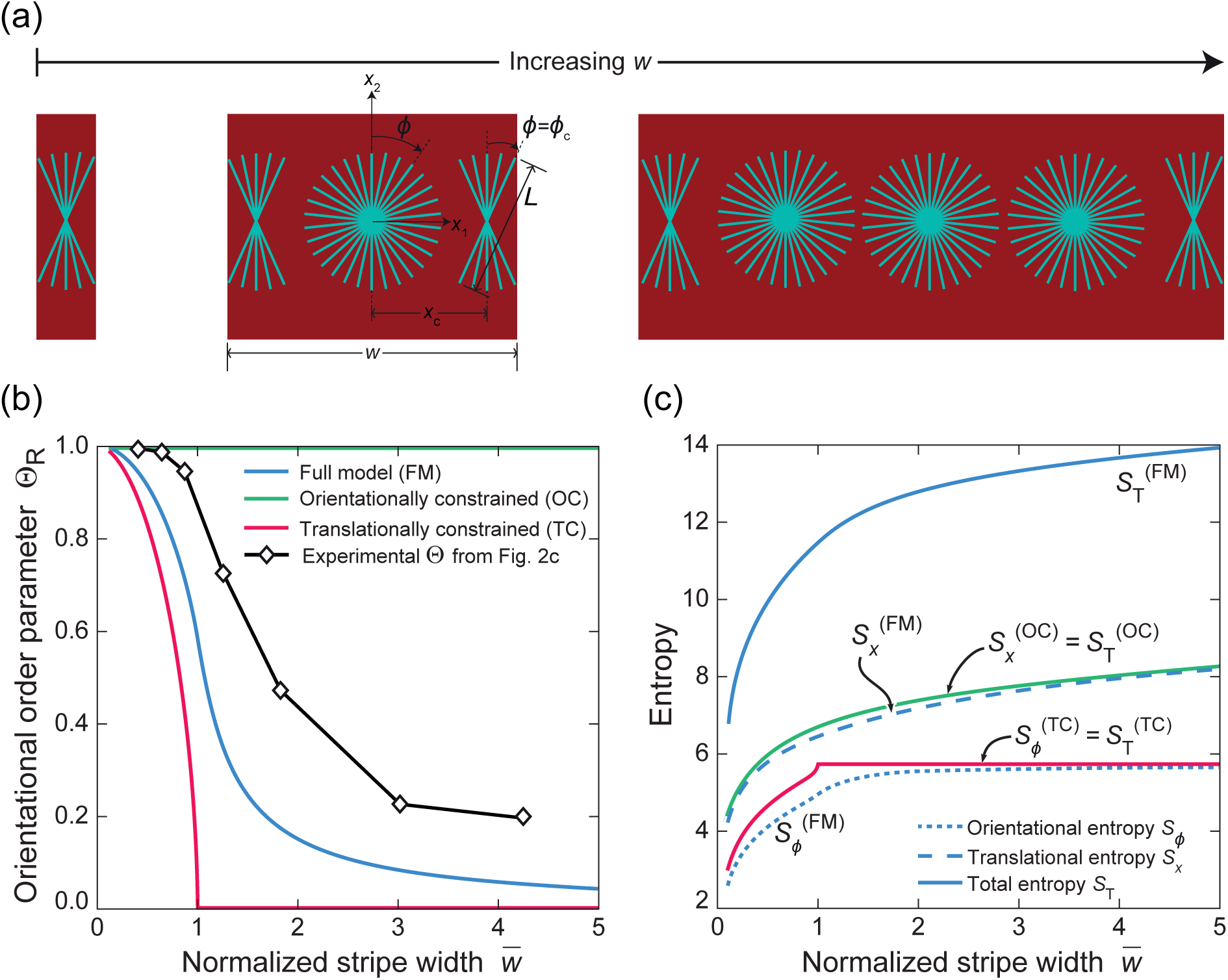
Entropic ordering of a hard-rod of length *L* in a channel of width *w*. (a) Sketch illustrating the states the hard-rod assumes as it translates and reorients within channels of three different normalised widths 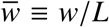. Rod orientation is denoted by φ. (b) Predictions of the orientational order parameter *Θ*_R_ as a function of 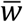 for the full model (blue) where the rod has translational and orientational degrees of freedom, the translationally constrained model (red) and the orientationally constrained model (green)), compared with the cell order parameter from experimental data on stripe widths normalised by the average cell length 2t_*e*_. (c) Corresponding predictions of the total entropy *S*_T_, orientational entropy *S*_φ_ and translational entropy *S_X_* for the three models as a function of 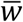. The discrete entropies plotted here are for the choices of the total number of available orientational and translational states *N*_φ_ = 314 and 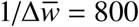, respectively (see Supplementary S4 for definitions).

To understand the preference of cells to follow a response consistent with the full model rather than a response in line with the translationally constrained model, we look at the corresponding entropies (Fig. 6c). We denote by 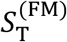 and 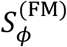 the total and orientational entropies, respectively, of the full model, while 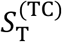 is the total entropy of the translationally constrained model (and equal to its orientational entropy 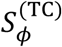. Clearly, 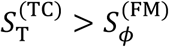 for all 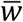 but 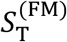 is significantly higher than 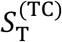, i.e., the full model has gained a higher total entropy by trading-off a small decrease in orientational entropy with a corresponding increase in its translational entropy. In other words, by allowing the rod to explore regions near the channel edges, one trades-off a small decrease in orientational entropy with a larger increase in translational entropy just as in the model of Onsager [34] for liquid crystals. Similarly, with 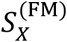 denoting the translational entropy of the full model and 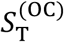 the total entropy of the orientationally constrained model (equal to its translational entropy 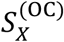, we observe that while 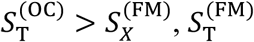 is significantly higher than 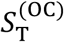.

In the experiments, fluctuations of the cell morphology are not orientationally or translationally restricted, with intracellular processes such as actin polymerisation and treadmilling capable of equally changing cell shape or enabling rotation and translation of the cell. Thus, fluctuation of cell morphology will result in a maximisation of morphological entropy with the cell assuming states akin to that for the full model, as seen from the experimental cell order parameter curve in Fig. 6b. The maximisation of morphological entropy results in partial ordering of specific observables and here, cell alignment emerges from the drive of cells to maximise their overall state of disorder. This simple hard rod model, while a powerful help to at-least qualitatively understand orientational ordering in regime I, has significant limitations as it does not encode any information on the cell biochemistry. As a consequence, it not only fails to account for changes in cell morphology in regime II but also has quantitative limitations in predicting the cell order parameter. The only parameter in the hard rod model is the rod length *L* and it is natural to equate that to the length 2*ℓ*_*e*_ = 130 μm of the major axis of the cells. The measurements of Θ with the normalised stripe width interpreted as 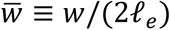 are included in Fig. 6b: clearly, while predictions of the hard rod model and the measurements have similar trends, the hard rod model significantly under-predicts the orientational ordering. The full homeostatic framework that includes critical biochemical processes such as the actomyosin stress-fibre remodelling (Fig. 1b) are indeed essential not only to understand the regimes of contact guidance but also to make quantitative predictions of the wide range of observables reported here.

## Discussion

Contact guidance, the phenomenon by which cell alignment is influenced by anisotropic patterns, not only has implications for how cells organise themselves into collective entities such as tissues, but is also critical in tissue engineering, where the geometric structure of scaffolds can be used to control the morphology and organisation of cells. Multiple hypotheses [8, 17, 18] have been proposed for rationalising the sensing mechanisms that drive contact guidance. While these theories highlight the role of specific cellular components including filopodia, focal adhesion, or stress-fibres, an understanding of why cells align with geometric or adhesive patterns has remained elusive.

Here, we use a statistical mechanics framework to interpret experimental data for the response of individual myofibroblasts on substrates with micropatterned stripes of fibronectin, and propose a novel explanation for contact guidance. This interpretation based on entropic considerations can be summarised as follows. The cell is an open system exchanging nutrients with its surrounding bath. Dynamic processes such as actin polymerisation and treadmilling are driven by biochemical reactions fuelled by these nutrients. These reactions are not precisely controlled and thus fluctuations in the morphological states of the cell ensue as widely recognised in the literature [35-37]. These cell shape fluctuations, occur on timescales longer than intracellular biochemical processes such as cytoskeletal remodelling and are ergodic in nature. This drives the cells towards a stationary state that is constrained by the fact that the coupled intracellular biochemical processes also strive to maintain the cell in a homeostatic state. These biochemical processes thus serve to provide both the mechanisms and constraints for the maximisation of morphological entropy. This enabled us to develop a statistical equilibrium formulation for the cell response and formalise the description of the non-thermal fluctuations through a homeostatic temperature 1/*ζ* that is conjugated to the morphological entropy. Our simulations show that the normalised homeostatic temperature of the myofibroblasts is on the order 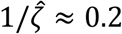(Fig. 5a), implying that 1/*ζ* is typically about 20% of the average Gibbs free-energy of all the morphological states the cell assumes. This translates to a homeostatic temperature 1/*ζ* ≈ 10^12^ K (Supplementary S2) clearly illustrating the non-thermal nature of the cell shape fluctuations. These large Gibbs free-energy fluctuations allow cells to change shape, rotate and wander over the fibronectin stripes so as to maximise their morphological entropy. However, the spatial constraint of the fibronectin stripes implies that cell morphologies close to the edges of the stripes need to be necessarily aligned with the edges of the stripes (Fig. 3). This results in an alignment of cells with the stripes that persists even when a representative cell size 2*ℓ*_e_ ≪ *w* and we refer to this as entropic contact guidance, labelled here as regime I. As the stripe width decreases to becomes of the order of 2*ℓ*_e_, the spatial constraints imposed by the fibronectin stripes induce biochemical changes within the cells that manifest as increases in the cell aspect ratio, greater polarisation of the stress-fibre arrangements within the cell, and a significantly enhanced level of cell alignment. We refer to this as the biochemically-mediated regime of contact guidance, labelled here as regime II.

The spatial constraint imposed by the fibronectin stripes on cells imposes guidance forces *F*_G_ that constrain cells to lie within the stripes and align the cells with the stripe direction. These forces comprise two components: (i) an entropic component *F*_E_ associated with the drive of the cell to maximise its morphological entropy, and (ii) an energetic (or biochemical) component *F*_B_ resulting from the drive of the cell to reduce the average Gibbs free-energy. It is thus natural to ask, what is the relation (if any) between the guidance force *F*_G_ and the traction forces exerted by cells on substrates? These traction forces are typically reported in experimental studies as measured via traction-force microscopy [38, 39], and can also be computed from our model. Predictions of normalised traction distributions 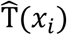 in randomly selected morphologies on three stripe widths are included in Fig. 5a along with the variation of the average normalised total traction force 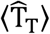 with *w* (see Supplementary S2.5.1 for detailed definitions). The average total traction force 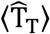 is invariant in regime I, but rises in regime II and clearly has a different functional dependence on stripe width compared with *F*_G_ shown in Fig. 5b. We defer a detailed discussion of this issue of the differences between 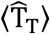 and *F*_G_ to Supplementary S3.2 but here it suffices to say that there is no direct relation between the forces exerted by cells on substrates via focal adhesions, which are of Newtonian nature, and the non-Newtonian guidance force *F*_G_. Indeed, *F*_G_ → 0 in the homogeneous substrate limit *w* → ∞, although the traction forces exerted by cells on the substrate do not vanish in this limit.

Current models of active matter typically either use the notion of an effective temperature that is empirically defined [36] or resort to non-equilibrium approaches such as hydrodynamic theories for active gels, where the concept of an affinity of energy carriers (e.g. ATP or GTP) is employed to maintain the system out of thermodynamic equilibrium [37]. The statistical framework employed here combines the reductionist approach of the hydrodynamic active matter theories with the simplicity of equilibrium formulations that employ an effective temperature. The key difference however lies in the fact that the homeostatic mechanics framework clearly defines the effective temperature within a statistical mechanics context, along with its conjugated entropy. The overall homeostatic mechanics framework is independent of the biochemical model used to describe the intracellular processes and the free-energy of a specific morphological microstate. Here, to model contact guidance we only considered contributions from the remodelling of the acto-myosin cytoskeleton, the passive elasticity of the cytoplasm as well as the nucleus along with adhesions that equilibrate forces generated by these cellular components (Fig. 1b). Thus, the kinetics of signalling pathways such as Rho signalling cascade triggered by the formation of focal adhesions and resulting in the polymerisation of stress-fibres are not explicitly modelled. Rather, in this equilibrium framework, the cytoskeletal structures, focal adhesion distributions as well as cell morphologies emerge as interlinked coupled phenomena. In fact, contact guidance is a direct outcome of the fact that near the stripe edges, the focal adhesion and cytoskeletal structures are restricted by the fibronectin patterns and this in turn imposes restrictions on the morphologies that are accessible over the course of the non-thermal fluctuations that the cells undergo.

In summary, we have shown that contact guidance of myofibroblasts by fibronectin stripes can be understood in terms of classical concepts of statistical thermodynamics, namely, entropy maximisation subject to appropriate constraints. In order to make these concepts applicable to the context of non-thermal morphological fluctuations of cells, we had to employ non-classical notions of entropy borrowed from information theory. This framework shows convincingly that contact guidance is based on physical forces of thermodynamic origin, due to shape fluctuations of the cell. The impact of this is multi-fold:

i. Since the fluctuations are of biochemical rather than of thermal origin, we do not need to invoke unrealistic thermal fluctuation energies to rationalise the behaviour of cells. The homeostatic mechanics framework formalises this notion of an effective temperature of cells by recognising that the fluctuations are related to the energy fluctuations resulting from nutrient exchanges with the nutrient bath.
ii. Our model predicts with remarkable fidelity not only trends but also the statistical variability of a wide range of biological observables. Indeed, the mechanism of contact guidance we propose here, viz. that contact guidance (or orientational ordering) emerges from the drive of cells to maximise the disorder of the morphological states they assume is inherently linked to this statistical variability.
iii. The concept of maximisation of morphological entropy subject to the constraint of homeostasis is very general and can be used to predict guidance as a function of cell phenotype as well as a range of biophysical and/or biochemical cues. For example, in Supplementary S2.6, we explore the question of whether the reduced stiffness of cancer cells [40] may result in enhanced contact guidance, thereby promoting tissue invasion via contact guidance [41, 42]. However, it remains to be seen via both experiments and corresponding predictions how widely the idea of maximisation of morphological entropy constrained by homeostasis is applicable to rationalise the response of cells to environmental cues more general than the ones examined in this study.

## Methods

Microcontact printing was used to pattern single (fibronectin) adhesive stripes of width *w* ranging from 50 µm to 1570 µm, spaced 500 µm apart using previously established protocols [43]. Briefly, the stripe patterns were generated on a silicon master by deep reactive-ion etching (Philips Innovation Services, Eindhoven, The Netherlands) from a chromium photomask (Toppan Photomask, Corbeil Essonnes, France). The silicon surface was passivated with fluorosilane and microstamps were obtained by moulding the silanized silicon master with polydimethylsiloxane (PDMS, Sylgard 184, Dow Corning) and a curing agent (10:1) and permitted to cure overnight at 65 °C. The cured PDMS stamp containing the desired features was then peeled off from the master and cleaned by sonicating it in 70% ethanol for 30 minutes and dried using compressed air. The structured surface of the PDMS stamps was inked for 1 hour at room temperature with a 50 mg ml^−1^ Rhodamine fibronectin (FN) solution (Cytoskeleton, Denver, CO, USA). Flat PDMS-coated glass coverslips served as the substrates on which microprinting was conducted. These substrates were first oxidized in a UV/Ozone cleaner (PDS UV-ozone cleaner; Novascan, Ames, IA) for 8 minutes, and then the FN-coated stamps (first dried under compressed air) were gently deposited on the substrates for 15 minutes at room temperature. Uncoated regions of the substrates were blocked by immersing the micropatterned coverslips for 5 minutes in a 1% solution of Pluronic F-127 (Sigma-Aldrich, St. Louis, MO). Finally, the coverslips were washed thrice with PBS and stored in PBS at 4 °C until use. A PDMS stamp without any features was used to print a homogeneous pattern of fibronectin on a flat PDMS surface, which was used as the control substrate.

### Cell culture

Human Vena Saphena Cells (HVSCs) were harvested from the vena saphena magna obtained from patients according to Dutch guidelines of secondary use material and have previously been characterized as myofibroblasts [44]. The myofibroblasts were cultured in Advanced Dulbecco’s Modified Eagle’s Medium (Invitrogen, Breda, The Netherlands) supplemented with 10% Fetal Bovine Serum (Greiner Bio-one), 1% penicillin/streptomycin (Lonza, Basel, Switzerland) and 1% GlutaMax (Invitrogen). Only cells with a passage lower than 7 were used in this study. To avoid cell-cell contact between the cells, the micropatterned substrates were seeded with a cell density of 500 cells cm^−2^ and cultured for 24 hours at 37 °C in 5% CO_2_.

### Cell fixation and staining

For visualization of the focal adhesions, actin cytoskeleton, and the nucleus, the myofibroblasts were fixed with 4% formaldehyde in PBS (Sigma-Aldrich) 24 hours after cell seeding. The cells were then permeabilized with 0.5% Triton-X-100 (Merck) for 15 minutes, blocked for 30 minutes with 4% goat serum in PBS, and incubated with the primary anti-vinculin antibody IgG1 (V9131, Sigma-Aldrich). Subsequently, samples were incubated with the secondary antibody goat anti mouse-Alexa Fluor 647 (Molecular Probes) 1:500 and FITC-conjugated phalloidin (15500, Phalloidin-Atto 488, Sigma-Aldrich) 1:200 for staining the actin cytoskeleton. Finally, the samples were incubated with DAPI (Sigma-Aldrich) for 5 minutes for immunofluorescence of the nucleus and mounted onto glass slides using Mowiol (Sigma-Aldrich). The images of the cells were acquired using an inverted microscope (Zeiss Axiovert 200M, Zeiss, Gottingen, Germany).

### Image analysis

Fluorescent microscopy images were recorded at 20 and 40 times magnification and then analysed using a custom-built Mathematica (Wolfram Research, Inc., Mathematica, Version 11.1) script as described previously [44]. The binarized images of the profile of each cell and nucleus were fitted with an ellipse using a least-square algorithm [44], and used to quantify observables that include: (i) orientation *φ* that measures the angle between the major axis of the best-fit ellipse and the stripe direction *x*_2_ (Fig. 2b); (ii) the aspect ratio *A*_s_ defined by the ratio of the lengths of the major to minor axis of the best-fit ellipse and (iii) the order parameter Θ defined as [21]

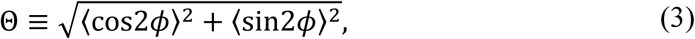

where 〈·〉 denotes the ensemble average over all measurements. The cell and nucleus area were directly determined from the area enclosed within the respective profiles; readers are referred to [44] for more details on the analysis of the data. For each stripe width, three independent experiments were performed and at least 50 cells were analysed per stripe width. Box-and-whisker diagrams (showing the quartiles, means and outliers) of cell and nuclear area, aspect ratio and orientation were constructed from the data for these 50 cells on each stripe width. The spatial distributions of the stress-fibre orientations 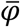 and the associated cytoskeletal order parameter ℛ were extracted from the actin-stained immunofluorescence images as described in detail in Supplementary S1.

## Supporting information

## Acknowledgements

ABCB acknowledges support from the TU/e Impuls and Gravitation grant. HS acknowledges support from the Commonwealth Scholarship Commission and Cambridge Trust. AV and VSD acknowledge support from the Royal Society’s Newton International Fellowship’s Alumni program. ADS gratefully acknowledges support from the European Research Council through ERC Advanced Grant 340685-Micromotility, and the warm hospitality of Corpus Christi College of the University of Cambridge.

## Footnotes

2 Similar to the fibronectin stripes that restrict the cell by preventing cell adhesion outside the stripes, we use the terminology of “channel” to indicate that the walls of the channel restrict the motion of a hard rod.

