## Supplementary material for "Entropic forces drive cellular contact guidance"

### Supplementary Information: Entropic forces drive cellular contact guidance

**Supplementary Information 1: Measurement of stress-fibre orientations**

The measurement of the distribution of stress-fibre orientations within the myofibroblasts is typically performed by the convolution of the so-called elongated Laplacian of Gaussian (eLoG) kernel with the image [1]. This procedure requires a choice of the anisotropic standard deviations used in the Gaussian kernel and they are chosen empirically to give good agreement between the predicted fibre orientations and those based on visual inspection. However, in our case, this procedure required modifications to the magnitudes of the standard deviations for every stripe width and thus, using this methodology risked introducing a bias in the measurements. The breakdown of this standard method in our case is related to the fact that unlike most previous studies, we were also interested in situations where stress-fibres were relatively randomly oriented and this required a higher resolution of stress-fibre detection in order to obtain adequate data. We thus followed a different method to infer the fibre orientations which involved first processing the images to highlight the stress-fibres and then analysing these pre-processed images.

Overall the procedure involved two steps to determine the spatial distribution $\varphi(x_i)$ of the actin stress-fibre orientations within the imaged cells: (i) image processing to highlight the actin fibres, and (ii) detection of fibres within these images and estimation of their orientations $\varphi$ with respect to the $x_2$ −direction of the fibronectin stripes. We shall now explain each of these two steps.

*1.1 Image processing to highlight the actin fibres*

Highlighting actin fibres is equivalent to edge detection in image processing and typically involves convolution of the image with a Laplacian kernel [2]. We did this in the following three steps (see also Fig. S4):

(a) The green channel from each of the immunofluorescence images (corresponding to the actin stains) was converted to a grayscale image and subsequently sharpened using the *imsharpen* function within MATLAB (standard deviation of the Gaussian low pass filter was set to 1, while 1.8 was used for the strength of the sharpening effect).

(b) The grayscale image from (a) was first smoothened by convolving with an isotropic $5 \times 5$ Gaussian kernel with a unit standard deviation and then convolved with the Laplacian kernel

$$L = \begin{bmatrix} 0 & -1 & 0 \\ -1 & 4 & -1 \\ 0 & -1 & 0 \end{bmatrix}, \quad (1.1)$$

to produce an image wherein the fibres are highlighted.

(c) Finally, the brightness of the image is adjusted by rescaling the pixel intensities $I$ to $I'$ via

$$I' = 1 + 45 \ln(I), \quad (1.2)$$

so as to brighten the edges (fibres) with a low intensity. This helps in reducing the fibre discontinuities that are artificially generated as a consequence of the convolutions in (b).

These operations produce a binary image with pixels of high intensity typically corresponding to (i) stress-fibres within the cell, or (ii) the perimeter of the cell. An example of the outcome of the image processing operations is shown in Fig. S4a starting from the green channel of a randomly selected immunofluorescence image of a myofibroblast on the homogeneous substrate, and followed by four images at each of the stages (a) through (c) described above.

#### 1.2 Fibre detection and estimation of orientation

Fibre detection was performed on the binary images with the highlighted fibres using an algorithm outlined in Obbink-Huizer et al. [3], which was first developed by Frangi et al. [4]. First the images were smoothened using a $5 \times 5$ Gaussian kernel with unit standard deviation and then edge detection was performed by calculating the Hessian matrix at each pixel using the Laplacian kernel (1.1). This Hessian was then used to define fibres and their orientation as described subsequently.

Denote the eigenvalues of the Hessian at pixel location $x_i$ to be $q_1(x_i)$ and $q_2(x_i)$ with $|q_1| \leq |q_2|$. The corresponding eigenvectors are $\boldsymbol{v}_1$ and $\boldsymbol{v}_2$. Now, recall that the ratio of eigenvalues $q_1/q_2$ is small for a pixel containing a stress-fibre since a fibre has a large magnitude of intensity gradient perpendicular to the fibre direction (i.e. large $|q_2|$ in direction $\boldsymbol{v}_2$), and a small magnitude of gradient parallel to the fibre direction (i.e. small $|q_1|$ in $\boldsymbol{v}_1$). Furthermore, fibres in the image have $q_2 < 0$ since the pixel intensity at these locations is higher compared to its surroundings. This understanding is used to define a measure of the likelihood that a pixel contains a stress-fibre. This measure is referred to as "fibreness" and defined as

$$\mathfrak{f} = \begin{cases} 0 & \text{if } \; q_2 \geq 0 \\ \exp\left[-2\left(\frac{q_1}{q_2}\right)^2\right] & \text{otherwise}. \end{cases} \tag{1.3}$$

The definition implies that $\mathfrak{f}$ attains a high value (defined here as $\mathfrak{f} \geq 0.6$) when the likelihood of the pixel being within a stress-fibre is high. All pixels with $\mathfrak{f} \geq 0.6$ (or equivalently $-q_2 \geq 2q_1$) are assumed to lie within a stress-fibre with the fibre oriented in direction $\boldsymbol{v}_1 = (v_1^x, v_1^y)$ where $v_1^x$ and $v_1^y$ are the components of $\boldsymbol{v}_1$ in the $x_1$ and $x_2$ directions, respectively. The orientation $\varphi(x_i)$ of the stress-fibre at location $x_i$ then follows as

$$\varphi = \tan^{-1}\left(\frac{v_1^y}{v_1^x}\right), \tag{1.4}$$

which is measured with respect to the stripe (i.e. $x_2$) direction. The stress-fibres detected using this algorithm are then coloured by their orientation $\varphi$ and shown in Fig. S4b, for the binary image from Fig. S4a.

In addition to using this procedure to visualise the stress-fibre distributions, we also performed statistical analysis of the data on the orientations of the stress-fibres generated by this procedure. The stress-fibre orientations $\varphi$ are not invariant to rigid body rotations of the cell. To interrogate the dispersion of stress-fibre orientations within the cell, we first defined a measure $\bar{\varphi}$ of the stress-fibre orientations that is invariant to rigid body rotations of the cell. An average stress-fibre orientation within an imaged cell is first calculated as

$$\langle \varphi \rangle = \frac{\sum \varphi \, \mathbb{I}(\mathfrak{f} \geq 0.6)}{\sum \mathbb{I}(\mathfrak{f} \geq 0.6)}, \tag{1.5}$$

where the summation is performed over all pixels in the image of the cell with $\mathbb{I}$ the indicator function. Then, $\bar{\varphi}(x_i) \equiv \varphi(x_i) - \langle \varphi \rangle$ is a rotationally invariant measure of stress-fibre orientation (i.e. $\langle \bar{\varphi} \rangle = 0$). The cell images in Fig. 4a (and Fig. S4b) show stress-fibres (inferred using the method described above) and coloured by their local orientations $\bar{\varphi}$. The distributions of $\bar{\varphi}$ over all imaged cells are then used to construct the probability density functions $p(\bar{\varphi})$ shown in Fig. 4c. The cytoskeletal orientational order parameter $\mathcal{R}$ is then calculated from $p(\bar{\varphi})$ via Eq. (1).

*1.3 Robustness of the definition of the cytoskeletal order parameter*

The cytoskeletal orientational order parameter is defined here over the ensemble of observations. In previous studies [1, 5], a different definition is employed: $\mathcal{R}$ is calculated using the distribution of stress-fibre orientations for a given cell morphology (i.e. for a given image of a cell), and then the mean and standard error of $\mathcal{R}$ are estimated over a number of imaged cells. We argue that the definition used here is not only more robust, but more appropriate when the response of cells is viewed in light of the homeostatic mechanics framework. This is clarified in Supplementary Section 2 so as to put the discussion in the context of the modelling framework.

**Supplementary Information 2: The homeostatic mechanics framework**

Here, we provide a brief overview of the homeostatic mechanics framework of Shishvan et al. [6]. The aim is to provide the reader the key aspects of the framework required for fully appreciating the computational results presented in the main text, and to also emphasise the differences in the present formulation compared to the work of Shishvan et al. [6]. Readers are referred to Shishvan et al. [6] for a more complete treatment including the derivations of the relevant equations.

Making the ansatz that *living cells are entropic,* Shishvan et al. [6] introduced the concept of the *homeostatic ensemble* with cellular homeostasis providing the additional constraints and mechanisms for entropy maximisation. This defined the notion of a (dynamic) *homeostatic equilibrium* state that intervenes to allow living cells to elude thermodynamic equilibrium. They thus developed a statistical mechanics framework for living cells using the notions of statistical inference [7] applicable over a timescale from a few hours to a few days as long as the cell remains as a single undivided entity (i.e. the interphase period of the cell cycle). The key ideas behind the framework can be summarised as follows. A system comprising the cell and the extracellular matrix (ECM) is an open system with the cell exchanging nutrients with the surrounding bath. These nutrients fuel a large number of coupled biochemical reactions that include actin polymerisation, treadmilling and dendritic nucleation that effect changes to the cell morphology. These biochemical reactions change the morphology of the cell but are not precisely controlled, and this manifests via the observed morphological fluctuations of the cell. Shishvan et al. [6] made the *ansatz* that these biochemical reactions provide the mechanisms to maximise the morphological entropy of the cell, but constrained by the fact that the cell maintains a homeostatic state[1] over the interphase period. Showing that the homeostatic constraint translates to a constraint on the average Gibbs free-energy, Shishvan et al. [6] developed a statistical mechanics framework to analyse the fluctuating response of cells. It is this framework that we extend to analyse contact guidance.

*2.1 Morphological microstates, entropy, fluctuations and the homeostatic temperature*

Controlling only macro variables (i.e. macrostate) such as the temperature, pressure and nutrient concentrations in the nutrient bath results in inherent uncertainty (referred to here as missing information) in micro variables (i.e. microstates) of the system. This includes a level of unpredictability in homeostatic process variables, such as the spatio-temporal distribution of chemical species, that is linked to Brownian motion and the complex feedback loops in the

[1] Cellular homeostasis is the ability of cells to actively regulate their internal state, and maintain the concentration of all internal species at specific average values over their morphological fluctuations independent of the environment.

homeostatic processes. Thus, this system not only includes the usual lack of precise information on the positions and velocities of individual molecules associated with the thermodynamic temperature, but also an uncertainty in cell shape resulting from imprecise regulation of the homeostatic processes. The consequent entropy production forms the basis of this new statistical mechanics framework motivated by the following two levels of microstates:
(i) *Molecular microstates*. Each molecular microstate has a specific configuration (position and momentum) of all molecules within the system.
(ii) *Morphological microstates (Fig. S5b)*. Each morphological microstate is specified by the mapping (connection) of material points on the cell membrane to material points on the substrate within the fibronectin stripes. In broad terms, a morphological microstate specifies the shape and size of the cell.

Shishvan et al. [6] identified the (dynamic) homeostatic or equilibrium state of the system by entropy maximisation. Subsequently, we shall simply refer to this state as an equilibrium state to emphasise that it is a stationary macrostate of the system inferred via entropy maximisation as in conventional equilibrium analysis. The total entropy of the system is written in terms of the conditional probability $P^{(i|j)}$ of the molecular microstate $(i)$ given the morphological microstate $(j)$ and the probability $P^{(j)}$ of morphological microstate $(j)$ as

$$I_{\mathrm{T}} = \sum_{j} P^{(j)} I_{\mathrm{M}}^{(j)} + I_{\Gamma}. \tag{2.1}$$

In Eq. (2.1), $I_{\mathrm{M}}^{(j)} \equiv -\sum_{i \in j} P^{(i|j)} \ln P^{(i|j)}$ and $I_{\Gamma} \equiv -\sum_{j} P^{(j)} \ln P^{(j)}$ are the entropies of molecular microstates in morphological microstate $(j)$ and the morphological microstates, respectively. Equilibrium then corresponds to molecular and morphological macrostates that maximise $I_{\mathrm{T}}$ subject to appropriate constraints. The molecular macrostate evolves on the order of seconds, limited by processes such as the diffusion of unbound actin. By contrast, transformation of the morphological macrostate involves cell shape changes and therefore, the morphological macrostate evolves on the order of minutes, limited by co-operative cytoskeletal processes within the cell such as meshwork actin polymerisation and dendritic nucleation. The evolutions of the molecular and morphological macrostates are therefore temporally decoupled, and Shishvan et al. [6] showed that Eq. (2.1) can be maximised by independently maximising $I_{\mathrm{M}}^{(j)}$ at the smaller timescale to determine the equilibrium distribution of molecular microstates for a given morphological microstate, and then maximising $I_{\Gamma}$ at the larger timescale to determine the equilibrium distribution of the morphological microstates.

Over the (short) timescale on the order of seconds, the only known constraint on the system is that it is maintained at a constant temperature, pressure and strain distribution. The equilibrium of a given morphological microstate $(j)$ obtained by maximising $I_{\mathrm{M}}^{(j)}$ (denoted by $S_{\mathrm{M}}^{(j)}$) corresponds to molecular arrangements that minimise the Gibbs free-energy with $G^{(j)}$. Since the connection between the cell and the substrate is fixed for a given morphological microstate, the determination of $G^{(j)}$ is a standard boundary value problem as described in Section 2.2. Over the (long) timescale on the order of several minutes to hours, the equilibrium distribution $P_{\mathrm{eq}}^{(j)}$ is determined by maximising $I_{\Gamma}$, but now with the additional constraint that the cell is maintained in its homeostatic state. For the case of a cell on an ECM in a constant temperature and pressure nutrient bath, the homeostatic constraint translates to the fact that the average Gibbs free-energy of the system over all the morphological microstates it assumes, is equal to the equilibrium Gibbs free-energy $G_{\mathrm{S}}$ of an isolated cell in suspension (free-standing cell), i.e. the homeostatic processes maintain the average biochemical state of the system equal to that of a cell in suspension. In deriving this result, Shishvan et al. [6] did not consider every individual homeostatic process, but rather used just the coarse-grained outcome of the homeostatic processes. The application of this coarse-grained constraint is the key element of the *homeostatic mechanics* framework, with the morphological entropy $I_\Gamma$ parameterising the information lost by not modelling all variables associated with the homeostatic processes.

The maximisation of $I_\Gamma$ while enforcing $\sum_j P^{(j)} G^{(j)} = G_S$ gives the *homeostatic equilibrium* state such that

$$P_{eq}^{(j)} = \frac{1}{Z} \exp\left(-\zeta G^{(j)}\right), \tag{2.2}$$

where $Z \equiv \sum_j \exp\left(-\zeta G^{(j)}\right)$ is the partition function of the morphological microstates, and the distribution parameter $\zeta$ follows from the homeostatic constraint

$$\frac{1}{Z} \sum_j G^{(j)} \exp\left(-\zeta G^{(j)}\right) = G_S \,. \tag{2.3}$$

The collection of all possible morphological microstates that the system assumes while maintaining its homeostatic equilibrium state is referred to as the *homeostatic ensemble*. The homeostatic ensemble can therefore be viewed as a large collection of copies of the system, each in one of the equilibrium morphological microstates. The copies $(j)$ are distributed in the ensemble such that the free-energies $G^{(j)}$ follow an exponential distribution $P_{eq}^{(j)}$ with the distribution parameter $\zeta$.

The equilibrium morphological entropy $S_\Gamma = -\sum_j P_{eq}^{(j)} \ln P_{eq}^{(j)}$ (i.e. the maximum value of $I_\Gamma$) follows from (2.2) and (2.3) as

$$S_\Gamma = \zeta G_S + \ln Z, \tag{2.4}$$

where $P_{eq}^{(j)}$ is substituted from Eq. (2.2). Thus, $S_\Gamma$ is related to $\zeta$ via the conjugate relation $\partial S_\Gamma / \partial G_S = \zeta$. Thus, analogous to $1/T$ that quantifies the increase in uncertainty of the molecular microstates (i.e. molecular entropy $S_M^{(j)}$) with average enthalpy, $\zeta$ specifies the increase in uncertainty of the morphological microstates (i.e. morphological entropy $S_\Gamma$) with the average Gibbs free-energy. We therefore refer to $1/\zeta$ as the *homeostatic temperature* with the understanding that it quantifies the fluctuations on a timescale much slower than that characterised by $T$.

#### *2.2 The equilibrium Gibbs free-energy of a morphological microstate*

Similar to conventional statistical mechanics calculations that require a model for the energy of the system, the homeostatic statistical mechanics framework requires a model for the Gibbs free-energy $G^{(j)}$ of morphological microstate $(j)$. Mathematical models of varying degrees of complexity [8-12] have been developed for cells subjected to specified boundary conditions and can be used to determine $G^{(j)}$. Here, we calculate $G^{(j)}$ using the free-energy model of Vigliotti et al. [9] that includes contributions from cell elasticity and the actin/myosin stress-fibre cytoskeleton, with the cell modelled as a two-dimensional (2D) body in the $x_1 - x_2$ plane adhered to a fibronectin stripe of width $w$ (Fig. S5c) such that the out-of-plane Cauchy stress $\Sigma_{33} = 0$. The state of the system changes as the cell moves, spreads and changes shape on the fibronectin stripe. Here, we shall give a prescription to calculate the Gibbs free-energy of the system when the cell is in a specific morphological microstate $(j)$, where the connections of material points on the cell membrane to the surface of the substrate are specified.

With the system comprising the cell and the rigid substrate within a constant temperature and pressure nutrient bath, the Gibbs free-energy $\mathcal{G}^{(j)}$ of the system in morphological microstate $(j)$ is given by $\mathcal{G}^{(j)} = \int_{V_{\text{cell}}} f\, dV$, where $f$ is the specific Helmholtz free-energy of the cell. Here, there is no contribution from the substrate as it is assumed to be rigid. We emphasize that the analysis presented here is for the system under atmospheric pressure conditions. Therefore, without loss of generality, we set the pressure equal to zero (i.e. use gauge pressure), and thus a pressure term does not appear in the expression for $\mathcal{G}^{(j)}$. The equilibrium free-energy $G^{(j)}$ is then the value of $\mathcal{G}^{(j)}$ at $d\mathcal{G}^{(j)} = 0$. Now, we briefly describe the model for the calculation of $G^{(j)}$. In the following, for the sake of notational brevity, we shall drop the superscript $(j)$ that denotes the morphological microstate, as the entire discussion refers to a single morphological microstate.

The Vigliotti et al. [9] model assumes only two elements within the cell: (i) a passive elastic contribution from elements such as the cell membrane, intermediate filaments and microtubules, and (ii) an active contribution from contractile acto-myosin stress-fibres that are modelled explicitly with the nucleus not explicitly modelled. This model was modified in [6] to incorporate a non-dilute concentration of stress-fibres and here we further modify this model by including the nucleus in the analysis as a passive elastic body, in addition to the cytoplasm comprising the two above mentioned components. We shall first describe the modelling of the active acto-myosin stress-fibres in the cytoplasm and then discuss the elastic model of both the nucleus and the cytoplasm.

Consider a two-dimensional (2D) cell of thickness $b_0$ and volume $V_0$ in its elastic resting state comprising a nucleus of volume $V_{\text{N}}$ and cytoplasm of volume $V_{\text{C}}$ such that $V_0 = V_{\text{N}} + V_{\text{C}}$ (Fig. S5c). The representative volume element (RVE) of the stress-fibres within the cytoplasm in this resting configuration is assumed to be a cylinder of volume $V_{\text{R}} = \pi b_0 \left(\frac{n^{\text{R}}\ell_0}{2}\right)^2$, where $\ell_0$ is the length of a stress-fibre functional unit in its ground-state, and $n^{\text{R}}$ is the number of these ground-state functional units within this reference RVE. The total number of functional unit packets within the cell is $N_0^{\text{T}}$, and we introduce $N_0 = N_0^{\text{T}} V_{\text{R}}/V_{\text{C}}$ as the average number of functional unit packets available per RVE; $N_0$ shall serve as a useful normalisation parameter. The state of stress-fibres at location $x_i$ within the cell is described by their angular concentration $\eta(x_i, \varphi)$, and there are $n(x_i, \varphi)$ functional units in series along the length of each stress-fibre in the RVE. Here, $\varphi$ is the angle of the stress-fibre bundle in the undeformed configuration with respect to the $x_2 -$ direction of the fibronectin stripe (Fig. S5c). Vigliotti et al. [9] showed that, at steady-state, the number $n^{\text{ss}}$ of functional units within the stress-fibres is given by

$$\hat{n}^{\text{ss}} \equiv \frac{n^{\text{ss}}}{n^{\text{R}}} = \frac{[1 + \varepsilon_{\text{nom}}(x_i, \varphi)]}{1 + \tilde{\varepsilon}_{\text{nom}}^{\text{ss}}}, \tag{2.5}$$

where $\tilde{\varepsilon}_{\text{nom}}^{\text{ss}}$ is the strain at steady-state within a functional unit of the stress-fibres, and $\varepsilon_{\text{nom}}(x_i, \varphi)$ is the nominal strain in direction $\varphi$. The chemical potential of the functional units within the stress-fibres in terms of the Boltzmann constant $k_{\text{B}}$ is given by [6]

$$\chi_{\text{b}} = \frac{\mu_{\text{b}}}{n^{\text{R}}} + k_{\text{B}} T \ln\left[\left(\frac{\pi\, \hat{\eta}\, \hat{n}^{\text{ss}}}{\hat{N}_{\text{u}}\left(1 - \frac{\hat{\eta}}{\hat{\eta}_{\max}}\right)}\right)^{\frac{1}{n^{\text{ss}}}} \left(\frac{\hat{N}_{\text{u}}}{\pi \hat{N}_{\text{L}}}\right)\right], \tag{2.6}$$

where the normalized concentration of the unbound stress-fibre proteins is $\hat{N}_{\text{u}} \equiv N_{\text{u}}/N_0$ with $\hat{\eta} \equiv \eta n^{\text{R}}/N_0$, while $\hat{\eta}_{\max}$ is the maximum normalised value of $\hat{\eta}$ corresponding to full occupancy of all available sites for stress-fibres (in a specific direction) and $\hat{N}_{\mathrm{L}}$ is the number of lattice sites available to unbound proteins. The enthalpy $\mu_{\mathrm{b}}$ of $n^{\mathrm{R}}$ bound functional units at steady-state is given in terms of the isometric stress-fibre stress $\sigma_{\max}$ and the internal energy $\mu_{\mathrm{b0}}$ as

$$\mu_{\mathrm{b}} = \mu_{\mathrm{b0}} - \sigma_{\max}\Omega(1 + \tilde{\varepsilon}_{\mathrm{nom}}^{\mathrm{ss}}), \tag{2.7}$$

where $\Omega$ is the volume of $n^{\mathrm{R}}$ functional units. By contrast, the chemical potential of the unbound proteins is independent of stress and given in terms of the internal energy $\mu_{\mathrm{u}}$ as

$$\chi_{\mathrm{u}} = \frac{\mu_{\mathrm{u}}}{n^{\mathrm{R}}} + k_{\mathrm{B}}T \ln\left(\frac{\hat{N}_{\mathrm{u}}}{\pi \, \hat{N}_{\mathrm{L}}}\right). \tag{2.8}$$

For a fixed configuration of the 2D cell (i.e. a fixed strain distribution $\varepsilon_{\mathrm{nom}}(x_i, \varphi)$), the contribution to the specific Helmholtz free-energy of the cell $f$ from the stress-fibre cytoskeleton follows as

$$f_{\mathrm{cyto}} = \rho_0 \left(\hat{N}_{\mathrm{u}}\chi_{\mathrm{u}} + \int_{-\pi/2}^{\pi/2} \hat{\eta} \, \hat{n}^{\mathrm{ss}} \chi_{\mathrm{b}} d\varphi\right), \tag{2.9}$$

where $\rho_0 \equiv N_0/V_{\mathrm{R}}$ is the number of protein packets per unit reference volume available to form functional units in the cell. However, we cannot yet evaluate $f_{\mathrm{cyto}}$ as $\hat{N}_{\mathrm{u}}(x_i)$ and $\hat{\eta}(x_i, \varphi)$ are unknown. These will follow from the chemical equilibrium of the cell as will be discussed in Section 2.2.1.

The total stress $\Sigma_{ij}$ within the cell includes contributions from the passive elasticity provided mainly by the intermediate filaments of the cytoskeleton attached to the nuclear and plasma membranes and the microtubules, as well as the active contractile stresses of the stress-fibres. The total Cauchy stress is written in an additive decomposition as

$$\Sigma_{ij} = \sigma_{ij} + \sigma_{ij}^{\mathrm{p}}, \tag{2.10}$$

where $\sigma_{ij}$ and $\sigma_{ij}^{\mathrm{p}}$ are the active and passive Cauchy stresses, respectively. In the 2D setting with the cell lying in the $x_1 - x_2$ plane, the active stress is given in terms of the volume fraction $\mathcal{H}_0$ of the stress-fibre proteins as

$$\begin{bmatrix} \sigma_{11} & \sigma_{12} \\ \sigma_{12} & \sigma_{22} \end{bmatrix} = \frac{\mathcal{H}_0\sigma_{\max}}{2} \int_{-\pi/2}^{\pi/2} \hat{\eta}[1 + \varepsilon_{\mathrm{nom}}(\varphi)] \begin{bmatrix} 2\sin^2\varphi^* & -\sin 2\varphi^* \\ -\sin 2\varphi^* & 2\cos^2\varphi^* \end{bmatrix} d\varphi, \tag{2.11}$$

where $\varphi^*$ is the angle of the stress-fibre measured with respect to $x_2$, and is related to its orientation $\varphi$ in the undeformed configuration by the rotation with respect to the undeformed configuration. The passive elasticity in the 2D setting is given by a 2D specialization of the Ogden [13] hyperelastic model as derived in [6]. The strain energy density function of this 2D Ogden model is

$$\Phi_{\mathrm{C}} \equiv \frac{2\mu_{\mathrm{C}}}{m_{\mathrm{C}}^2}\left[\left(\frac{\lambda_{\mathrm{I}}}{\lambda_{\mathrm{II}}}\right)^{\frac{m_{\mathrm{C}}}{2}} + \left(\frac{\lambda_{\mathrm{II}}}{\lambda_{\mathrm{I}}}\right)^{\frac{m_{\mathrm{C}}}{2}} - 2\right] + \frac{\kappa_{\mathrm{C}}}{2}(\lambda_{\mathrm{I}}\lambda_{\mathrm{II}} - 1)^2, \tag{2.12}$$

for the cytoplasm and

$$\Phi_{\mathrm{N}} \equiv \frac{2\mu_{\mathrm{N}}}{m_{\mathrm{N}}^2}\left[\left(\frac{\lambda_{\mathrm{I}}}{\lambda_{\mathrm{II}}}\right)^{\frac{m_{\mathrm{N}}}{2}} + \left(\frac{\lambda_{\mathrm{II}}}{\lambda_{\mathrm{I}}}\right)^{\frac{m_{\mathrm{N}}}{2}} - 2\right] + \frac{\kappa_{\mathrm{N}}}{2}(\lambda_{\mathrm{I}}\lambda_{\mathrm{II}} - 1)^2, \tag{2.13}$$

for the nucleus where $\lambda_{\mathrm{I}}$ and $\lambda_{\mathrm{II}}$ are the principal stretches, $\mu_{\mathrm{C}}$ ($\mu_{\mathrm{N}}$) and $\kappa_{\mathrm{C}}$ ($\kappa_{\mathrm{N}}$) the shear modulus and in-plane bulk modulus of cytoplasm (nucleus), respectively, while $m_{\mathrm{C}}$ ($m_{\mathrm{N}}$) is a material constant governing the non-linearity of the deviatoric elastic response of cytoplasm (nucleus). The cell is assumed to be incompressible, and thus throughout the cell, we set the principal stretch in the $x_3$ −direction $\lambda_{\mathrm{III}} = 1/(\lambda_{\mathrm{I}}\lambda_{\mathrm{II}})$. The (passive) Cauchy stress then follows as $\sigma_{ij}^{\mathrm{p}} p_j^{(k)} = \sigma_k^{\mathrm{p}} p_i^{(k)}$ in terms of the principal (passive) Cauchy stresses $\sigma_k^{\mathrm{p}}$ ($\equiv \lambda_k \partial\Phi_{\mathrm{C}}/\partial\lambda_k$ for the cytoplasm and $\equiv \lambda_k \partial\Phi_{\mathrm{N}}/\partial\lambda_k$ for the nucleus) and the unit vectors $p_j^{(k)}$ ($k = \mathrm{I}, \mathrm{II}$) denoting the principal directions. The total specific Helmholtz free-energy of the cell is then $f = f_{\mathrm{cyto}} + \Phi_{\mathrm{C}}$ in the cytoplasm and $f = \Phi_{\mathrm{N}}$ in the nucleus.

##### 2.2.1 *Equilibrium of the morphological microstate*

Shishvan et al. [6] have shown that equilibrium of a morphological microstate reduces to two conditions: (i) mechanical equilibrium with $\Sigma_{ij,j} = 0$ throughout the system, and (ii) chemical equilibrium such that $\chi_{\mathrm{u}}(x_i) = \chi_{\mathrm{b}}(x_i, \varphi) = \mathrm{constant}$, i.e. the chemical potentials of bound and unbound stress-fibre proteins are equal throughout the cell. The condition $\chi_{\mathrm{u}} = \chi_{\mathrm{b}}$ implies that $\hat{\eta}(x_i, \varphi)$ is given in terms of $\hat{N}_{\mathrm{u}}$ by

$$\hat{\eta}(x_i, \varphi) = \frac{\hat{N}_{\mathrm{u}}\, \hat{\eta}_{\max} \exp\left[\frac{\hat{n}^{\mathrm{ss}}(\mu_{\mathrm{u}} - \mu_{\mathrm{b}})}{k_{\mathrm{B}}T}\right]}{\pi \hat{n}^{\mathrm{ss}} \hat{\eta}_{\max} + \hat{N}_{\mathrm{u}} \exp\left[\frac{\hat{n}^{\mathrm{ss}}(\mu_{\mathrm{u}} - \mu_{\mathrm{b}})}{k_{\mathrm{B}}T}\right]}, \tag{2.14}$$

and $\hat{N}_{\mathrm{u}}$ follows from the conservation of stress-fibre proteins throughout the cytoplasm, viz.

$$\hat{N}_{\mathrm{u}} + \frac{1}{V_{\mathrm{C}}} \int_{V_{\mathrm{C}}} \int_{-\pi/2}^{\pi/2} \hat{\eta}\, \hat{n}^{\mathrm{ss}} d\varphi \ dV = 1. \tag{2.15}$$

Knowing $\hat{N}_{\mathrm{u}}$ and $\hat{\eta}(x_i, \varphi)$, the stress $\Sigma_{ij}$ can now be evaluated and these stresses within the system (i.e. cell and substrate) need to satisfy mechanical equilibrium, i.e. $\Sigma_{ij,j} = 0$. In this case, the mechanical equilibrium condition is readily satisfied as the stress field $\Sigma_{ij}$ within the cell is equilibrated by a traction field $\mathrm{T}_i$ exerted by the substrate on the cell such that $b\Sigma_{ij,j} = -\mathrm{T}_i$, where $b(x_i)$ is the thickness of the cell in the current configuration.

The equilibrium value of $\mathcal{G}$ denoted by $G$ is then given as $G = F_{\mathrm{cell}}$ where

$$F_{\mathrm{cell}} \equiv \rho_0 V_{\mathrm{C}} \chi_{\mathrm{u}} + \int_{V_{\mathrm{C}}} \Phi_{\mathrm{C}}\, dV + \int_{V_{\mathrm{N}}} \Phi_{\mathrm{N}}\, dV. \tag{2.16}$$

Here, $\chi_{\mathrm{u}}$ is given by Eq. (2.8) with the equilibrium value of $\hat{N}_{\mathrm{u}}$ obtained from Eq. (2.15). For the purposes of further discussion, we label the equilibrium value $F_{\mathrm{cyto}} \equiv \rho_0 V_{\mathrm{C}} \chi_{\mathrm{u}}$ as the cytoskeletal free-energy of the cell, and $F_{\mathrm{passive}} \equiv \int_{V_{\mathrm{C}}} \Phi_{\mathrm{C}}\, dV + \int_{V_{\mathrm{N}}} \Phi_{\mathrm{N}}\, dV$ as the passive elastic energy of the cell.

#### 2.3 *Numerical methods*

We employ Markov Chain Monte Carlo (MCMC) to construct a Markov chain that is representative of the homeostatic ensemble. This involves three steps: (i) a discretization scheme to represent morphological microstate ($j$), (ii) calculation of $G^{(j)}$ for a given morphological microstate ($j$), and (iii) construction of a Markov chain comprising these morphological microstates. Here, we briefly describe the procedure which was implemented in MATLAB with readers referred to [6] for further details.

In the general setting of a three-dimensional (3D) cell, a morphological microstate is defined by the connection of material points on the cell membrane to the surface of the substrate. In the 2D context of cells on micropatterned substrates, this reduces to specifying the connection of all material points of the cell to locations within the fibronectin stripes, i.e. a displacement field $u_i^{(j)}(X_i)$ is imposed on the cell with $X_i$ denoting the location of material points on the cell in the undeformed configuration, and these are then displaced to $x_i^{(j)} = X_i + u_i^{(j)}$ in morphological microstate $(j)$, such that all $x_i^{(j)}$ lie within the fibronectin stripe. These material points located at $x_i^{(j)}$ are then connected to material points on the substrate at the same location $x_i^{(j)}$, completing the definition of the morphological microstate in the 2D setting.

The cell is modelled as a continuum and thus $u_i^{(j)}$ is a continuous field. To calculate the density of the morphological microstates, we define $u_i^{(j)}$ via Non-Uniform Rational B-splines (NURBS) such that the morphological microstate is now defined by $M$ pairs of weights $\left[U_L^{(j)}, V_L^{(j)}\right]$ $(L = 1, \dots, M)$. In all the numerical results presented here, we employ $M = 16$ with $4 \times 4$ weights $U_L^{(j)}$ and $V_L^{(j)}$ governing the displacements in the $x_1$ and $x_2$ directions, respectively. The NURBS employ fourth order base functions for both the $x_1$ and $x_2$ directions, and the knots vector included two nodes each with multiplicity four, located at the extrema of the interval. We emphasise here that this choice of representing the morphological microstates imposes restrictions on the morphological microstates that will be considered. Therefore, the choice of the discretization used to represent $u_i^{(j)}$ needs to be chosen so as to be able to represent the microstates we wish to sample, e.g. the choice can be based on the minimum width of a filopodium one expects for the given cell type. Given $u_i^{(j)}$, we can calculate $G^{(j)}$ using the model described in Section 2.2 with the cell discretised using constant strain triangles of size $e \approx R_0/10$, where $R_0$ is the radius of the cell in its undeformed configuration.

We construct, via MCMC, a Markov chain that serves as a sample of the homeostatic ensemble for cells on fibronectin stripes of width $w$. The algorithm closely follows the approach developed by Shishvan et al. [6]. However, here we are modelling cells constrained on fibronectin stripes with cell adhesion outside the stripes prevented. Over the range of stripe widths used in the experimental investigation, cells that were partially or entirely outside the fibronectin stripes were not observed. Thus, we construct a sample of the homeostatic ensemble comprising solely of morphological microstates that are fully within the fibronectin stripes. This is done using the Metropolis [14] algorithm in an iterative manner using the procedure explained in detail in [6] (see *section 4.3* therein) but now with the following modification. In constructing the Markov chain, if any portion of the proposed new configuration of the cell lies outside the fibronectin stripe, then the nodal boundary points outside the fibronectin stripe were pushed back to the boundary of the fibronectin stripe along a line joining their locations in the current proposed morphological microstate and their corresponding positions in previous accepted morphological microstate- this corrected morphological state was then used as the configuration on which to check the probability of acceptance. Typical Markov chains comprised in excess of $\mathcal{L} = 4 \times 10^6$ samples.

##### *2.4 Material parameters for myofibroblasts*

All simulations are reported at a reference thermodynamic temperature $T = T_0$, where $T_0 = 310$ K. Most of the parameters of the model are related to the properties of the proteins that constitute stress-fibres. These parameters are thus expected to be independent of cell type.

Notable exceptions to this are: (i) the stress-fibre protein volume fraction $\mathcal{H}_0$; and (ii) the passive elastic properties. Here, we use parameters calibrated for myofibroblasts that give good correspondence with the wide range of measurements reported here. The passive elastic parameters of the cytoplasm are taken to be $\mu_C = 1.7$ kPa, $\kappa_C = 35$ kPa and $m_C = 5$, while the corresponding values for the nucleus are $\mu_N = 3.4$ kPa, $\kappa_N = 35$ kPa and $m_N = 20$. The maximum contractile stress $\sigma_{max} = 240$ kPa is consistent with a wide range of measurements on muscle fibres [15], and the density of stress-fibre proteins was taken as $\rho_0 = 3 \times 10^6$ μm$^{-3}$ with the volume fraction of stress-fibre proteins $\mathcal{H}_0 = 0.032$. Following Vigliotti et al. [9], we assume that the steady-state functional unit strain $\tilde{\varepsilon}_{nom}^{ss} = 0.35$ with $\mu_{b0} - \mu_u = 2.3\ k_B T_0$ and $\Omega = 10^{-7.1}$ μm$^3$. The maximum angular concentration of stress-fibre proteins is set to $\hat{\eta}_{max} = 0.75$. The cell in its undeformed state is a circle of radius $R_0$ and thickness $b_0$, with a circular nucleus of radius $R_N$ whose centre coincides with that of the cell. The radius of the myofibroblasts in their undeformed state was taken to be $R_0 = 30$ μm, while their thickness was set at $b_0/R_0 = 0.2$. In order to estimate $R_N$, the myofibroblasts on the homogenous substrate were treated with blebbistatin. This inhibited the actomyosin network forcing the cells to adopt their undeformed state. The cell and nucleus diameter were measured to infer that the nucleus occupies a volume fraction $\bar{v}_N = 0.21$ of the cell. Within the two-dimensional setting of this numerical study, this implies $R_N/R_0 = \sqrt{\bar{v}_N} = 0.45$.

*2.5 Definitions of normalised quantities and observables*

Following [6], the free-energy $G^{(j)}$ can be decomposed as $G^{(j)} = \Upsilon^{(j)} + \Upsilon_0$, where $\Upsilon_0 \equiv \rho_0 V_0 \left[\mu_u/n^R - k_B T \ln\left(\pi\, \hat{N}_L\right)\right]$ is independent of the morphological microstate. It is thus natural to subtract out $\Upsilon_0$ and define a normalised free-energy as

$$\hat{G}^{(j)} \equiv \frac{\Upsilon^{(j)}}{|G_S - \Upsilon_0|} = \frac{G^{(j)} - \Upsilon_0}{|G_S - \Upsilon_0|}\ , \tag{2.17}$$

where $G_S$ is the equilibrium free-energy of a free-standing cell (i.e. a cell in suspension with traction-free surfaces). Then, the distribution given by Eq. (2.2) can be re-written as

$$P_{eq}^{(j)} = \frac{1}{\hat{Z}} \exp\left[-\hat{\zeta} \hat{G}^{(j)}\right], \tag{2.18}$$

with $\hat{Z} \equiv \sum_j \exp\left[-\hat{\zeta}\hat{G}^{(j)}\right]$ and $\hat{\zeta} \equiv \zeta |G_S - \Upsilon_0|$. It then immediately follows that the distributions of states are not influenced by the values of $n^R$, $\hat{N}_L$ and $V_0$ and these parameters need not be specified so long as energies are quoted in terms of the normalised energies $\hat{G}^{(j)}$. We note in passing that we have normalised the Gibbs free-energy of each morphological microstate and the homeostatic temperature by the homeostatic value of the Gibbs free-energy that the cell attains over all the morphological microstates it samples. For the myofibroblast parameters listed in Section 2.4, the cell in suspension has a radius of $\approx 27$ μm with a Gibbs free-energy $G_S - \Upsilon_0 \approx -7.7 \times 10^{10} k_B T_0 = -0.3$ nJ (see Shishvan et al. [6] for details of the calculation of $G_S - \Upsilon_0$). This implies that in units of Kelvin, a typical normalised temperature $1/\hat{\zeta} = 0.2$ of myofibroblasts (Fig. 5a) corresponds to $1/\zeta \approx 10^{12}$ K. Of course, this high temperature stems from the homeostatic framework inherently recognising that the morphological fluctuations of cells, rather than having a thermal origin, are biochemical in nature and arise from the imprecise regulation of the exchange of nutrients with the nutrient bath.

Analogously, we define the normalised passive and cytoskeletal free-energies of the cell as

$$\hat{F}_{passive}^{(j)} \equiv \frac{F_{passive}^{(j)}}{|G_S - \Upsilon_0|}, \tag{2.19}$$

and

$$\hat{F}_{\text{cyto}}^{(j)} \equiv \frac{F_{\text{cyto}}^{(j)} - \Upsilon_0}{|G_S - \Upsilon_0|}, \tag{2.20}$$

respectively. Probability density functions showing predictions of the distributions of $\hat{G}$, and $\hat{F}_{\text{cyto}}$ for myofibroblasts on selected widths of fibronectin stripes are included in Figs. S3a and S3b, respectively. These distributions are generated by plotting histograms from the sample list generated via the MCMC procedure and normalising the frequencies to give $p(\text{x})$ such that $\int_{-\infty}^{\infty} p(\text{x})d\text{x} = 1$, where the dummy symbol x denotes either $\hat{G}$ or $\hat{F}_{\text{cyto}}$. In line with the regimes of contact guidance discussed in the main text, the energy distributions are invariant to stripe width $w$ in regime I, but with biochemical changes being induced within the cell in regime II ($w \leq 160$ μm), the cytoskeletal free-energy $\hat{F}_{\text{cyto}}$ becomes more negative in regime II with decreasing $w$. This is a direct result of the aspect ratio of the cell increasing which causes additional stress-fibre polymerisation and the consequent reduction in $\hat{F}_{\text{cyto}}$. The reduction in $\hat{F}_{\text{cyto}}$ also directly influences the distribution of $\hat{G}$ which can be understood as follows. A reduction in $\hat{F}_{\text{cyto}}$ has a tendency to induce a higher probability of states with a lower $\hat{G}$. However, the homeostatic constraint enforces the ensemble average value of $\hat{G}$ to be fixed at $\hat{G} = -1$ and thus, the cell also increasingly samples states with high $\hat{G}$. This is typically achieved by increased straining of the cell resulting in increased values of $\hat{F}_{\text{passive}}$ (Fig. S3c). The overall consequence is a wider distribution of $\hat{G}$ within the homeostatic ensemble for myofibroblasts in regime II, accompanied by an increase in homeostatic temperature $1/\hat{\zeta}$ (Fig. 5a).

##### 2.5.1 *Tractions exerted by cells on substrates*

An observable commonly reported in experiments, typically measured via traction-force microscopy, are spatial distributions of tractions exerted by cells on substrates and the associated so-called total traction force [16-18]. These measurements are nearly exclusively reported on relatively soft substrates where tractions exerted by the cell induce significant deformations. This makes measurements of surface displacements feasible and allows for a relatively low error estimate of the associated surface tractions [17, 18]. Nevertheless, it is well-established that the response of cells is mechano-sensitive in the sense that the tractions they exert depend on the substrate stiffness, with the tractions increasing with increasing substrate stiffness [19]. The experiments in this study were conducted on relatively stiff substrates making measurements of tractions impractical, but predictions of tractions can be extracted from the simulations. Mechanical equilibrium (Section 2.2.1) of a given morphological microstate $(j)$ specifies the traction distribution $\text{T}^{(j)}(x_i)$ between the cell and the substrate. We define a normalised resultant traction

$$\hat{\text{T}}^{(j)}(x_i) \equiv \frac{R_0\sqrt{\text{T}_1^2 + \text{T}_2^2}}{b_0\mu_C}, \tag{2.21}$$

where $R_0$ and $b_0$ are the radius and thickness of the cell in its undeformed state, respectively, $\mu_C$ is the shear modulus of the cytoplasm and $T_i$ is the traction at location $x_i$ in morphological state $(j)$. These distributions for selected morphological microstates on three widths of fibronectin stripes are included in Fig. 5a. In addition, we also define the normalised total traction force as

$$\hat{\text{T}}_{\text{T}}^{(j)} \equiv \frac{1}{A^{(j)}} \int_{A^{(j)}} \hat{\text{T}}^{(j)} dA, \tag{2.22}$$

and the average value $\langle \widehat{\mathrm{T}}_{\mathrm{T}} \rangle$ of the total tractions (obtained as an average of the generated Markov chain) is included in Fig. 5a as a function of the fibronectin stripe width.

##### *2.5.2 Construction of immunofluorescence-like images*

The predictions in Fig. 1c which show immunofluorescence-like images comprise four layers; fibronectin stripes are depicted in maroon, the nucleus is coloured in blue while the focal adhesions and actin stress-fibres in shades of pink and green, respectively. Focal adhesion distributions in the experiments were observed by staining for vinculin (Fig. 1b). In the current set of simulations, the focal adhesions were not explicitly modelled but rather the cell was assumed to perfectly adhere to the substrate without directly accounting for the distribution of adhesion proteins. This results in the simulations predicting traction distributions $\mathrm{T}_i(x_i)$ as discussed above. However, it is well-known that traction magnitudes scale with concentration of adhesive proteins [20] and thus, here we use predictions of traction magnitudes as a surrogate to visualise the predictions of focal adhesion distributions. Specifically, we include in Fig. 1c distributions of $\widehat{\mathrm{T}}^{(j)}$ for selected morphological microstates $(j)$ with the darker shades of pink representing higher values of $\widehat{\mathrm{T}}^{(j)}$ and thereby higher vinculin concentrations

The predictions of the actin stress-fibre structure within the cytoplasm are coloured in green. These images, are meant to give two key pieces of information: (i) the local orientation of the dominant stress-fibre bundle, and (ii) the concentration of the stress-fibre proteins. However, no discrete fibres are present in the model of the acto-myosin stress-fibres employed here with the fibres represented via continuum internal state variables. Here, we use this continuum information to generate a discrete depiction of the fibres using the following prescription. Recall that for the purposes of calculation of $G^{(j)}$, the cell was discretized into 800 constant strain triangles of approximately equal size in the undeformed configuration. Within each element $k$, we defined an actin stress-fibre concentration in direction $\varphi^*$ as $\hat{\vartheta}_{\mathrm{b}}^{k}(\varphi^*) \equiv \hat{\eta}(\varphi^*)\hat{n}^{\mathrm{ss}}(\varphi^*)$, where $\varphi^*$ is the angle of the stress-fibre measured with respect to $x_2$ direction and is related to its angle $\varphi$ in the undeformed configuration by the rotation of element $k$. Next, we construct a dummy mesh comprising approximately 200 triangular elements to discretize the deformed cell. Elements of the original computational mesh are assigned to the element of the dummy mesh in which their centroid lies. In this manner, approximately four elements of the original computational mesh are assigned to each element of the dummy mesh. We then define an average actin stress-fibre concentration for each element $m$ of the dummy mesh as

$$\langle \hat{\vartheta}_{\mathrm{b}}(\varphi^*) \rangle_m \equiv \frac{1}{V_m} \sum_{k \in m} \hat{\vartheta}_{\mathrm{b}}^{k}(\varphi^*) v_k \, , \tag{2.23}$$

where $v_k$ is the volume of element $k$ of the computational mesh that is assigned to element $m$ of the dummy mesh and $V_m = \sum_{k\in m} v_k$. The dominant stress-fibre direction $\varphi^*_{\max}$ in dummy element $m$ is defined as the value of $\varphi^*$ corresponding to the maximum value of $\langle \hat{\vartheta}_{\mathrm{b}}(\varphi^*) \rangle_m$. Each element of the dummy mesh is hatched with green lines in the direction $\varphi^*_{\max}$. We set the spacing of the hatches to scale with the local density of bound stress-fibre proteins in order to replicate the higher intensity of the fluorescent phalloidin observed under light microscopy in locations with a higher concentration of stress-fibres. The total concentration of bound stress-fibre proteins in element $k$ of the computational mesh is

$$\widehat{N}_{\mathrm{b}}^{k} = \int_{-\pi/2}^{\pi/2} \hat{\vartheta}_{\mathrm{b}}^{k}(\varphi) \, d\varphi \, , \tag{2.24}$$

with $\widehat{N}_{\mathrm{b}}^{\mathrm{T}} \equiv \int_{V_{\mathrm{C}}} \widehat{N}_{\mathrm{b}} \, dV$ denoting the total concentration of bound stress-fibre proteins in the cell.

The spacing $\mathcal{s}$ of the green hatches in element $m$ of the dummy mesh is set to scale inversely with the concentration of bound stress-fibre proteins within that element, i.e. $\mathcal{s} \equiv \hat{N}_{\mathrm{b}}^{T} / \sum_{k \in m} \hat{N}_{\mathrm{b}}^{k}$ . This process results in the predictions of the actin cytoskeletal structure shown in Fig. 1c, and we include in Fig. S6 enlarged versions of some of those images along with insets showing zoom-ins to give more insight into the details of the predicted cytoskeletal structure at various locations within the cell.

##### *2.5.3 Distributions of stress-fibre orientations*

Given the spatial distributions $\varphi_{\max}^{*}(x_i)$, we followed a procedure analogous to the experimental protocol elaborated in Supplementary S1 to define a rotationally invariant spatial distribution of stress-fibre orientations. Specifically, we defined the rotationally invariant measure of stress-fibre orientation as

$$\bar{\varphi} \equiv \varphi_{\max}^{*} - \frac{1}{V_{\mathrm{C}}} \int_{V_{\mathrm{C}}} \varphi_{\max}^{*} \, dV \, , \tag{2.25}$$

with the predictions of $\bar{\varphi}(x_i)$ included in Fig. 4b. Consistent with observations, micro-domains of aligned stress-fibres emerge from the calculations (e.g. portions of the cell reminiscent of a filopodia have aligned stress-fibres in one orientation but misaligned with other parts of the cell). Moreover, similar to the observations, the selected images shown in Fig. 4b suggest a wider distribution of $\bar{\varphi}$ on wider stripes compared to the narrower stripes, and we wish to quantify the distribution of $\bar{\varphi}$ over the entire homeostatic ensemble. Recall that the computational mesh comprises 800 constant strain triangles, of which approximately 640 discretize the cytoplasm. Thus, for every morphological microstate generated by the MCMC, we have 640 stress-fibre orientations $\bar{\varphi}$ and a list of $640\mathcal{L}$ orientations over the entire sample of the homeostatic ensemble. Since all the elements in the undeformed state have approximately equal volume and the cytoplasm is incompressible, each of these orientations is over approximately the same volume of the cell. We therefore assigned equal weightages to each of the $640\mathcal{L}$ orientations and binned these orientations into intervals of size $\Delta\bar{\varphi} = \pi/4095$. The resulting histogram was then used to construct a probability density function $p(\bar{\varphi})$. Predictions of $p(\bar{\varphi})$ are included in Fig. 4c for selected fibronectin stripe widths and are in excellent agreement with the corresponding measurements.

##### *2.5.4 The cytoskeletal orientational order parameter*

The cytoskeletal orientational order parameter $\mathcal{R}$ is defined here over the ensemble of observations and calculated using Eq. (1), and the probability density functions $p(\bar{\varphi})$ from Fig. 4c. This ensures that the definition of $\mathcal{R}$ used in the predictions is consistent with the definition used in the experimental study (Supplementary S1), and these predictions are included in Fig. 4d. However, $\mathcal{R}$ as defined here differs from that employed in previous experimental studies [1, 5], and it is worth elaborating on the differences and the reasons for our choice.

In previous experimental studies [1, 5], the cytoskeletal orientational order parameter is calculated using the distribution of stress-fibre orientations for a given cell morphology (i.e. for a given cell image). For the sake of clarity, we shall label this as $\mathcal{R}^{(j)}$ to emphasize that this is an order parameter for a given morphological microstate $(j)$. The mean $\langle \mathcal{R}^{(j)} \rangle$ and standard error of $\mathcal{R}^{(j)}$ are then reported based on a finite number of measured values $\mathcal{R}^{(j)}$. Within the context of the homeostatic mechanics framework, each morphological microstate is random but the ensemble of these microstates is constrained to attain a state of homeostasis. This constraint sets the probability of observing a particular morphological microstate. The distribution of stress-fibre orientations within a specific morphological microstate is completely set by the specification of the microstate. Thus, the cytoskeletal orientational order parameter $\mathcal{R}^{(j)}$ for each morphological microstate $(j)$ is random but trends will emerge when the statistics of the parameter are investigated over the entire homeostatic ensemble. However, given that $\mathcal{R}^{(j)}$ is already a statistical measure of the stress-fibre orientations, interrogating the statistics of this statistical measure over the homeostatic ensemble (i.e. statistics of a statistic) is expected to lead to large errors. To explain this in a quantitative manner, recall that for a given probability density function $p(\bar{\varphi})$, the orientational order parameter $\mathcal{R}$ is the circular variance of $p(\bar{\varphi})$. Analytical expressions for required quantities are available for the variance of distributions, and thus, we proceed to explain the difficulties with using the usual literature definition of the orientational order parameter by using the variance of $p(\bar{\varphi})$ as a surrogate for the order parameter.

Consider the distribution $p(\bar{\varphi})$ over all states the cell assumes in a given environment (in our case a given width of fibronectin stripes) with

$$a_n \equiv \int_{-\frac{\pi}{2}}^{\frac{\pi}{2}} [\bar{\varphi} - \langle\bar{\varphi}\rangle]^n \, p(\bar{\varphi}) d\bar{\varphi}, \tag{2.26}$$

denoting the $n^{\text{th}}$ central moment of $p(\bar{\varphi})$, and $\langle\bar{\varphi}\rangle$ the mean of $\bar{\varphi}$ (the definition of $\bar{\varphi}$ implies that $\langle\bar{\varphi}\rangle = 0$). Within the context of the homeostatic mechanics framework, each morphological microstate (or imaged cell) comprises a distribution of stress-fibre orientations drawn randomly from $p(\bar{\varphi})$. To measure the variance $b_2^{(j)}$ of stress-fibre orientations within a morphological microstate $(j)$ (here $b_2^{(j)}$ serves as a surrogate for $\mathcal{R}^{(j)}$), we randomly draw $\mathcal{N}$ samples from $p(\bar{\varphi})$, i.e. we assume that each morphological microstate comprises $\mathcal{N}$ independent and identically distributed (*i.i.d.*) random orientations. The expected value of $b_2^{(j)}$ (over a large number of independent measurements of $b_2^{(j)}$) is then given by [21]

$$\langle b_2^{(j)} \rangle = \frac{\mathcal{N}-1}{\mathcal{N}} a_2. \tag{2.27}$$

Thus, in line with our intuition, $\langle b_2^{(j)} \rangle$ scales with the variance of $p(\bar{\varphi})$. Now consider the expected variance of $b_2^{(j)}$ in order to get a measure of the spread in the measured values of $b_2^{(j)}$. Kenney and Keeping [21] show that

$$\langle \text{var}(b_2^{(j)}) \rangle = \frac{(\mathcal{N}-1)^2}{\mathcal{N}^3} a_4 - \frac{(\mathcal{N}-1)(\mathcal{N}-3)}{\mathcal{N}^3} a_2^2, \tag{2.28}$$

which implies that the variance of the individual measurements $b_2^{(j)}$ depends both on the variance $a_2$ and kurtosis $a_4$ of $p(\bar{\varphi})$. With the kurtosis being a measure of the tailedness [22] of $p(\bar{\varphi})$, we anticipate a large variability in $b_2^{(j)}$ over the homeostatic ensemble.

In fact, in our preliminary simulations, we attempted to define the cytoskeletal order parameter for each individual morphological microstate within the Markov chain analogous to the experimental protocols reported in [1, 5]. However, we faced precisely the problem alluded to above, viz. $\mathcal{R}^{(j)}$ within the Markov chain had a very large dispersion and this made it impossible to determine the statistics of $\mathcal{R}^{(j)}$ with any confidence. Based on the above discussion, this is not surprising from the perspective of the model. However, what is more interesting is the fact that a large variability in $\mathcal{R}^{(j)}$ is also evident in reported measurements; see for example *Fig. 4* in Gupta et al. [5]. This suggests that the experimental system (cells) is behaving in a manner similar to the assumptions within the homeostatic mechanics framework, i.e. each morphological state is random with constraints (viz. homeostasis) only present over the ensemble of morphological states the cell samples. Therefore, both from a physical perspective as well as from considerations of mathematical robustness, we favour the definition of $\mathcal{R}$ employed in this study, with $\mathcal{R}$ directly calculated from $p(\bar{\varphi})$ over the ensemble of observations.

*2.6 Cell elasticity and contact guidance*

It has been argued that the reduced stiffness of cancer cells [23] is related to their enhanced ability for tissue invasion via contact guidance [24, 25]. Given the fidelity of the model in capturing a wide range of observations for myofibroblasts on fibronectin stripes, here we interrogate the model to understand the relation between cell stiffness (elasticity) and contact guidance. In order to explore the role of elasticity in terms of a single scalar variable, we restrict our analysis to the case where we vary the shear moduli of the cytoplasm and nucleus such that $\hat{\mu}_{\mathrm{C}} = \varsigma\mu_{\mathrm{C}}$ and $\hat{\mu}_{\mathrm{N}} = \varsigma\mu_{\mathrm{N}}$, where $\mu_{\mathrm{C}}$ and $\mu_{\mathrm{N}}$ are the corresponding moduli for the reference case of myofibroblasts, and keep the exponents $m_{\mathrm{C}}$ and $m_{\mathrm{N}}$ fixed at the values for myofibroblasts. Thus, all subsequent results are presented in terms of the scalar $\varsigma$ that quantifies the change in the cell elasticity with respect to myofibroblasts such that $\varsigma = 1$ for myofibroblasts.

Predictions of the cell order parameter $\Theta$ are included in Fig. S7a for two cases of stiffer cells in addition to the reference case of myofibroblasts with $\varsigma = 1$. Clearly, increasing stiffness decreases alignment (or equivalently, contact guidance) for any given stripe width. This reduction in contact guidance is related to two effects of increasing the cell stiffness:

(i) Modification of the distribution of shapes that the cells assume over their homeostatic state.

(ii) Reduction in the homeostatic temperature.

We shall understand the effect of both these changes in turn. The probability density functions of the cell area and aspect ratio are included in Figs. S7b and S7c, respectively for cells on homogenous substrates ($w \to \infty$). With increasing $\varsigma$, both the mean area $\bar{A}$ and mean aspect ratio $\bar{A}_{\mathrm{s}}$ decrease, with a consequent reduction in the leading cell length $2\ell_{\mathrm{e}} = 2\sqrt{(\bar{A}_{\mathrm{s}}\bar{A})/\pi}$. Recalling that the stripe width to transition from regime I to regime II scales approximately as $w_{\mathrm{crit}} \approx 2\ell_{\mathrm{e}}$, increasing cell stiffness implies that the transition from entropic to the stronger, biochemically-mediated contact guidance occurs for narrower stripe widths. However, this does not explain the lower levels of contact guidance over the entire range of stripe widths.

To understand these generally lower levels of contact guidance for stiffer cells, we include in Fig. S7d predictions of the variation of the normalised homeostatic temperature $1/\hat{\zeta}$ with $w$: for all $\varsigma$, $1/\hat{\zeta}$ increases with decreasing $w$, but more importantly, $1/\hat{\zeta}$ is lower for the stiffer cells for all $w$. The decrease in $1/\hat{\zeta}$ with increasing cell stiffness can be understood by recalling that the homeostatic temperature quantifies fluctuations in the Gibbs free-energy of morphological microstates, driven by the fact that cell biochemistry is not precisely regulated. Thus, increasing cell stiffness and hence the passive elastic contribution to the cell free-energy reduces the influence of these biochemical fluctuations and results in the consequent reduction of the homeostatic temperature. Concurrent with the decrease in homeostatic temperature, not only do the mean values of cell shape metrics such as area and aspect ratio decrease, but the variance in the distributions of these metrics also decreases; see Figs. S7b and S7c. Thus, we would expect fewer morphological microstates of the stiffer cells to interact with the stripe edges leading to a reduction in the strength of contact guidance. This is essentially an entropic argument and consistent with the fact that the entropic component of the guidance force, which dominates over the entire range of stripe widths, scales as $F_{\mathrm{E}} \propto 1/\hat{\zeta}$ (see Eq. (3.11) in Supplementary S3).

**Supplementary Information 3: The thermodynamic guidance force**

The inability of cells to adhere to the substrate outside the fibronectin stripes constrains the cells to stay within the stripes. Thus, similar to a gas within a container whose volume is constrained by the pressure exerted by the container on the gas (and vice-versa), the fibronectin stripes are expected to generate a guiding force that constrains cells to lie within the stripes. We shall use the homeostatic statistical mechanics framework to precisely define this guidance force, and thereby also develop an understanding of the elements that contribute to it.

Consider a cell at homeostatic equilibrium within a fibronectin stripe of width $w$. The morphological microstates of the cell fluctuate over the homeostatic ensemble such that the ensemble average Gibbs free-energy of the system comprising the cell and extracellular matrix (ECM) is given as

$$G_{\mathrm{S}} = \bar{G} \equiv \sum_j P^{(j)} G^{(j)}, \tag{3.1}$$

where, for the sake of brevity, we have reduced the notation of the equilibrium probability of a morphological microstate $(j)$ to read as $P^{(j)}$ rather than $P_{\mathrm{eq}}^{(j)}$. In line with the usual notion of thermodynamic forces, the guidance force $F_{\mathrm{G}}$ will be defined based on variations $d\bar{G}$ in the average Gibbs free-energy. An infinitesimal variation $dw$ in the fibronectin stripe width results in a change in $\bar{G}$ due to (i) a change in the probability $P^{(j)}$ of the energy levels; (ii) changes to $G^{(j)}$, and (iii) changes in the average number $\bar{N}_\alpha$ of each of the molecular species $(\alpha)$ within the cell such that

$$d\bar{G} = \sum_j \left(\frac{\partial \bar{G}}{\partial P^{(j)}}\right)_{G^{(j)},\bar{N}_\alpha} dP^{(j)} + \left(\frac{\partial \bar{G}}{\partial w}\right)_{P^{(j)},\bar{N}_\alpha} dw + \sum_{(\alpha)} \chi_\alpha d\bar{N}_\alpha. \tag{3.2}$$

Here, $\chi_\alpha$ is the chemical potential of species $(\alpha)$ in the homeostatic ensemble and the summation $\sum_{(\alpha)}(\cdot)$ is over all species $(\alpha)$ in the system. The average number $\bar{N}_\alpha$ of each these species is defined in terms of their number $N_\alpha^{(j)}$ in each morphological microstate via $\bar{N}_\alpha \equiv \sum_j P^{(j)} N_\alpha^{(j)}$. Then, upon using the definition (3.1), it follows

$$d\bar{G} = \sum_j G^{(j)} dP^{(j)} + \left(\frac{\partial \bar{G}}{\partial w}\right)_{P^{(j)},\bar{N}_\alpha} dw + \sum_{(\alpha)} \chi_\alpha d\bar{N}_\alpha. \tag{3.3}$$

We shall now proceed to show that the first term on the right-hand side of Eq. (3.3) is related to the morphological entropy. Recall that the morphological entropy $S_\Gamma$ is only a function of $P^{(j)}$ such that

$$dS_\Gamma = \sum_j \frac{\partial S_\Gamma}{\partial P^{(j)}} dP^{(j)}, \tag{3.4}$$

with $S_\Gamma = -\sum_j P^{(j)} \ln P^{(j)}$. It then follows that $dS_\Gamma = -\sum_j \ln P^{(j)} dP^{(j)}$, and substituting for $P^{(j)}$ from (2.2) we have

$$dS_\Gamma = \zeta \sum_j G^{(j)} dP^{(j)}. \tag{3.5}$$

Substituting back in (3.3), we obtain

$$d\bar{G} = \frac{dS_\Gamma}{\zeta} + \left(\frac{\partial \bar{G}}{\partial w}\right)_{S_\Gamma, \bar{N}_\alpha} dw + \sum_{(\alpha)} \chi_\alpha d\bar{N}_\alpha, \tag{3.6}$$

where we have noted that keeping $P^{(j)}$ constant is equivalent to a constant morphological entropy $S_\Gamma$. We then define the thermodynamic guidance force as $F_\text{G} \equiv (\partial \bar{G}/\partial w)_{S_\Gamma, \bar{N}_\alpha}$ and then analogous to the common statement of the first law of thermodynamics, we have an expression for the guidance force as

$$F_\text{G} dw = d\bar{G} - \frac{dS_\Gamma}{\zeta} - \sum_{(\alpha)} \chi_\alpha d\bar{N}_\alpha. \tag{3.7}$$

At homeostatic equilibrium, while the Gibbs free-energy $G^{(j)}$ of the cell fluctuates, the homeostatic potential $\mathcal{M} \equiv -(1/\zeta)\ln Z$ is a constant [6]. Recalling Eq. (2.4), i.e. $S_\Gamma = \zeta\bar{G} + \ln Z$, it follows that $\mathcal{M} = \bar{G} - (1/\zeta)S_\Gamma$. Thus, completely analogous to the usual definition of pressure in conventional thermodynamics, the guidance force is given by

$$F_\text{G} = \left(\frac{\partial \mathcal{M}}{\partial w}\right)_{\zeta, \bar{N}_\alpha} = \left(\frac{\partial \bar{G}}{\partial w}\right)_{S_\Gamma, \bar{N}_\alpha}, \tag{3.8}$$

with $\mathcal{M}$ taking the place of the Helmholtz free-energy, and $\bar{G}$ playing the role of internal energy.

It is instructive to write the guidance force as

$$F_\text{G} = \left(\frac{\partial \mathcal{M}}{\partial w}\right)_{\zeta, \bar{N}_\alpha} = \left(\frac{\partial \bar{G}}{\partial w}\right)_{\zeta, \bar{N}_\alpha} - \frac{1}{\zeta}\left(\frac{\partial S_\Gamma}{\partial w}\right)_{\zeta, \bar{N}_\alpha}, \tag{3.9}$$

where we emphasise that $\bar{G}$ and $S_\Gamma$ are now not fundamental thermodynamic functions[2]. The guidance force written in the form (3.9) enables us to distinguish the biochemical and entropic contributions as follows. Recall that the Gibbs free-energy $G^{(j)}$ of the cell is set by the elastic deformations and cytoskeletal arrangements within the cell. Thus, the first term $(\partial \bar{G}/\partial w)_{\zeta, \bar{N}_\alpha}$, which denotes the change in the average Gibbs free-energy of the cell, is a measure of the change in the cell biochemistry due to a change in the width of the fibronectin stripes. On the other hand, the second term of the right-hand side of Eq. (3.9) is the entropic force resulting from fluctuations of the morphological microstates over the homeostatic ensemble with the magnitudes of these fluctuations set by the homeostatic temperature $1/\zeta$. With homeostatic equilibrium being attained at a minimum value of $\mathcal{M}$, the biochemical force, given by

$$F_\text{B} \equiv \left(\frac{\partial \bar{G}}{\partial w}\right)_{\zeta, \bar{N}_\alpha}, \tag{3.10}$$

originates from the cell attempting to minimise its average Gibbs free-energy, while the entropic force, given by

$$F_\text{E} \equiv -\frac{1}{\zeta}\left(\frac{\partial S_\Gamma}{\partial w}\right)_{\zeta, \bar{N}_\alpha}, \tag{3.11}$$

originates from the cell's tendency to maximise its morphological entropy $S_\Gamma$. The total guidance force is $F_\text{G} = F_\text{B} + F_\text{E}$.

---

[2] The fundamental thermodynamic potentials written in terms of their natural variables are $\bar{G}(S_\Gamma, w, \bar{N}_\alpha)$ and $S_\Gamma(\bar{G}, w, \bar{N}_\alpha)$ with extremums of these potentials providing the equilibrium state. However, $\bar{G}(\zeta, w, \bar{N}_\alpha)$ and $S_\Gamma(\zeta, w, \bar{N}_\alpha)$ in (3.9) are only components of the fundamental potential $\mathcal{M}(\zeta, w, \bar{N}_\alpha)$ and individually will not predict the state of equilibrium. Nevertheless, these non-fundamental potentials, analogous to internal energy written as a function of thermodynamic temperature while calculating the Helmholtz free-energy, provide useful tools.

##### 3.1 Numerical methods

The numerical method used to evaluate these thermodynamic forces from the MCMC calculations requires some special considerations that are described here. For the purposes of numerical evaluation, the total guidance force is most conveniently expressed through the partition function as

$$F_{\mathrm{G}} = -\frac{1}{\zeta}\left(\frac{\partial \ln Z}{\partial w}\right)_{\zeta,\bar{N}_\alpha} = -\frac{1}{Z\zeta}\left(\frac{\partial Z}{\partial w}\right)_{\zeta,\bar{N}_\alpha}, \tag{3.12}$$

where the partition function written in terms of a continuous integral is given by

$$Z = \frac{1}{\Lambda^{2M} M!}\int \exp[-\zeta G(\boldsymbol{r}^M)]\, d\boldsymbol{r}^M\ . \tag{3.13}$$

Here, $\boldsymbol{r}$ denotes the pairs of weights that set a morphological microstate, and $\Lambda^{2M}$ is the "volume" of phase space that the $M$ pairs of weights occupy (i.e. analogous to the de Broglie wavelength, $\Lambda$ sets the minimum discrete unit by which the weights can vary). Defining a set of weights $\boldsymbol{s} \equiv \boldsymbol{r}/w$ scaled by the linear dimension of the system, it follows

$$Z = \frac{w^M}{\Lambda^{2M} M!}\int \exp[-\zeta G(\boldsymbol{s}^M)]\, d\boldsymbol{s}^M\ . \tag{3.14}$$

The domain of the integral in Eq. (3.14) is now independent of $w$. This considerably simplifies the calculation of the gradient with respect to $w$ and $F_{\mathrm{G}}$ readily follows from Eq. (3.12) as

$$F_{\mathrm{G}} = -\frac{M}{\zeta w} + \frac{1}{Z}\int \left[\frac{\partial G(\boldsymbol{s}^M)}{\partial w}\right] \exp[-\zeta G(\boldsymbol{s}^M)]\, d\boldsymbol{s}^M = -\frac{M}{\zeta w} + \langle \frac{\partial G}{\partial w} \rangle. \tag{3.15}$$

Here $\langle\cdot\rangle$ denotes the average over the homeostatic ensemble, and the calculation of $F_{\mathrm{G}}$ for a given stripe width $w$ therefore reduces to the calculation of $\langle \partial G/\partial w\rangle$. This ensemble average is computed as follows. Consider a cell on a substrate with a fibronectin stripe width $w$ in morphological microstate $(j)$ with Gibbs free-energy $G^{(j)}$. The substrate is subjected to a strain increment $\delta\varepsilon_{11}$ in the $x_1$ −direction causing a variation $\delta w = w\delta\varepsilon_{11}$ in the width of the fibronectin stripe. With the cell adhered to the stripe, its morphology is assumed to change affinely, and the variation in its Gibbs free-energy follows as

$$\frac{\partial G^{(j)}}{\partial w} = \frac{1}{w}\frac{\partial G^{(j)}}{\partial \varepsilon_{11}}\ . \tag{3.16}$$

For each morphological microstate in the Markov chain generated by the Metropolis sampling, perturbed cell morphologies are generated by applying the affine transformation $\Delta x_1 = \varepsilon_{11} x_1$ to all material points on the cell. These perturbed states are used to evaluate $\partial G^{(j)}/\partial\varepsilon_{11}$ via an adaptive central difference scheme with Romberg extrapolation used to generate error estimates for the adaptive scheme. The gradient $\partial G^{(j)}/\partial w$ is then calculated from Eq. (3.16), and the ensemble average $\langle \partial G^{(j)}/\partial w\rangle$ over the Markov chain is used as the estimate of $\langle \partial G/\partial w\rangle$.

Calculation of the biochemical force $F_{\mathrm{B}}$ is made tractable by noting that

$$F_{\mathrm{B}} \equiv \left(\frac{\partial \bar{G}}{\partial w}\right)_{\zeta,\bar{N}_\alpha} = \left(\frac{\partial \bar{G}}{\partial \zeta}\right)_{w,\bar{N}_\alpha}\left(\frac{\partial \zeta}{\partial w}\right)_{\bar{G},\bar{N}_\alpha}. \tag{3.17}$$

We first discuss the evaluation of $(\partial\bar{G}/\partial\zeta)_{w,\bar{N}_\alpha}$. Recall that

$$\bar{G} = \frac{w^M}{\Lambda^{2M} M!}\int G(\boldsymbol{s}^M)\exp[-\zeta G(\boldsymbol{s}^M)]\, d\boldsymbol{s}^M, \tag{3.18}$$

and differentiating (3.18) it follows that

$$\left[\frac{\partial \bar{G}}{\partial(1/\zeta)}\right]_{w,\bar{N}_\alpha} = \zeta^2[\langle G^2\rangle - \bar{G}^2]\,. \tag{3.19}$$

In analogy with the specific heat capacity of a solid at constant volume, $(\partial\bar{G}/\partial\zeta)_{w,\bar{N}_\alpha}$ is proportional to the variance in the Gibbs free-energy of the cell (i.e. the Gibbs free-energy fluctuation). The biochemical guidance force is then given as

$$F_{\mathrm{B}}(w) \equiv \zeta^2[\langle G^2\rangle - \bar{G}^2]\left[\frac{\partial(1/\zeta)}{\partial w}\right]_{\bar{G},\bar{N}_\alpha}. \tag{3.20}$$

There are three terms in the above expression: (i) $\zeta^2$ is given directly from the known homeostatic temperature of the particular line width (Fig. 5a); (ii) the variance of the Gibbs free-energy for a given stripe width follows from the variance of the Gibbs free-energy within the Markov chain generated in the numerical calculations for the stripe width under consideration at the correct homeostatic temperature, and (iii) the gradient of the homeostatic temperature $[(\partial(1/\zeta)/\partial w]_{\bar{G},\bar{N}_\alpha}$. This gradient needs to be calculated such that $\bar{G}$ and $\bar{N}_\alpha$ are held fixed while varying the stripe width. However, recall that the definition of homeostasis implies $\bar{N}_\alpha$ is constant, and Shishvan et al. [6] demonstrated that (for a given cell type) this also implies a constant $\bar{G}$. Thus, homeostatic temperature calculated for all stripe widths necessarily requires that $\bar{G}$ and $\bar{N}_\alpha$ are equal for all the stripe widths, and therefore both $\bar{G}$ and $\bar{N}_\alpha$ are constant along the curve of $1/\zeta$ versus $w$ shown in Fig. 5a. Thus, $[(\partial(1/\zeta)/\partial w]_{\bar{G},\bar{N}_\alpha}$ follows directly from the gradient of this curve.

In summary, the guidance force is given by

$$F_{\mathrm{G}} = F_{\mathrm{B}} + F_{\mathrm{E}} = \zeta^2[\langle G^2\rangle - \bar{G}^2]\left[\frac{\partial(1/\zeta)}{\partial w}\right]_{\bar{G},\bar{N}_\alpha} - \frac{1}{\zeta}\left(\frac{\partial S_\Gamma}{\partial w}\right)_{\zeta,\bar{N}_\alpha}, \tag{3.21}$$

with the entropic force calculated as the residual force $F_{\mathrm{E}} = F_{\mathrm{G}} - F_{\mathrm{B}}$.

##### *3.2 The nature of the guidance forces*

Prior to discussing the nature of the guidance forces described above, it is first worth briefly reviewing forces in classical thermodynamics (e.g. the canonical ensemble). This will enable us to easily clarify the distinction between those classical forces and the forces within the homeostatic ensemble. Consider a system comprising $N$ particles and maintained at temperature $T$ and volume $V$, and thus described by the canonical ensemble. At equilibrium, the Helmholtz potential $\mathcal{F} \equiv \bar{U} - TS$ is constant in this ensemble, with $\bar{U}$ and $S$ denoting the thermodynamic internal energy and entropy, respectively. Then, the thermodynamic force associated with a change in the volume of the system follows as

$$F = \left(\frac{\partial \mathcal{F}}{\partial V}\right)_{T,N} \equiv C_v\left(\frac{\partial T}{\partial V}\right)_{\bar{U},N} - T\left(\frac{\partial S}{\partial V}\right)_{T,N}, \tag{3.22}$$

where, in terms of the Boltzmann constant $k_{\mathrm{B}}$,

$$C_v \equiv \left(\frac{\partial \bar{U}}{\partial T}\right)_{V,N} = \frac{1}{k_{\mathrm{B}}T^2}\left[\langle U^2\rangle - \bar{U}^2\right] \tag{3.23}$$

is the specific heat capacity at constant volume. Here, $U$ denotes the internal energy of a particular microstate, while $\bar{U} \equiv \langle U\rangle$ is the ensemble average of the internal energies over the canonical ensemble, and $\langle U^2\rangle - \bar{U}^2$ the corresponding variance. The energetic and entropic components of the total force are $C_v(\partial T/\partial V)_{\bar{U},N}$ and $-T(\partial S/\partial V)_{T,N}$, respectively, and it is evident that both these forces are directly related to the thermodynamic temperature $T$. A microstate within the canonical ensemble is specified through the canonical co-ordinates of position and momentum $(\boldsymbol{q},\boldsymbol{p})$, and the temporal evolution of these co-ordinates is governed by a Hamiltonian consistent with Newton's laws of motion. The temperature $T$ sets the magnitude of the fluctuations of this Newtonian Hamiltonian, and thus the force in (3.22) is Newtonian, e.g. the pressure of a gas at temperature $T$ and constrained within a container of volume $V$.

It is evident that there are many similarities between (3.22) combined with (3.23), and the corresponding expression (3.21) for the guidance force within the homeostatic ensemble. In fact, the two expressions are identical with the following terms reinterpreted: (i) the internal energy is replaced by the Gibbs free-energy; (ii) the thermodynamic temperature is replaced by the homeostatic temperature, and (iii) consistent with the appropriate temperature conjugates, the molecular entropy $S$ is replaced by the morphological entropy $S_{\Gamma}$. Thus, we can interpret the guidance forces within the homeostatic ensemble in a manner analogous to the forces in the context of the canonical ensemble. In particular, the biochemical force within the homeostatic ensemble is given by the product of the specific Gibbs free-energy capacity and the gradient of the homeostatic temperature (3.20), while the entropic force is the product of the homeostatic temperature and the gradient of the morphological entropy, i.e. Eq. (3.11). However, while there exist these formal similarities between forces in the traditional thermodynamic ensembles and the homeostatic ensemble, there are critical quantitative and physical differences between the forces. Similar to forces in the canonical ensemble, forces in the homeostatic ensemble are directly related to the homeostatic temperature. However, unlike the canonical ensemble, the morphological microstates within the homeostatic ensemble are only specified by the positions of material points of the cell (which in turn specify the cell morphology), with no reference made to a co-ordinate for the velocities of these material points. In fact, the homeostatic ensemble is constructed using the ideas of statistical inference theory without recourse to any underlying Hamiltonian that specifies the evolution of the morphological microstates. This is one of the strengths of the homeostatic mechanics framework as it circumvents the issue that *the Hamiltonian for cells is unknown*. However, the consequent drawback is that the homeostatic temperature can only be interpreted as a distribution constant that sets the uncertainty or variability in the morphological microstates so as to maximise the morphological entropy while satisfying the necessary constraints on the system, i.e. we cannot assign a physical interpretation to the homeostatic temperature such as the average kinetic energy of the cell. Thus, the guidance force (3.21), although having the units of force, is *not* a Newtonian force. We prefer to think of the thermodynamic forces associated with the homeostatic ensemble, much like free-energies in traditional thermodynamics, as mathematical constructs that cannot be directly measured but useful to help interpret the behaviour of cells.

#### Supplementary Information 4: Entropic alignment of a hard rod in a channel

To illustrate the entropic alignment of cells constrained to lie within fibronectin stripes of width $w$, we consider the simple case of a hard rod (one-dimensional) of length $L$ within a channel of width $w$. The microstates of the rod are described in terms of the position $(x_1, x_2) = (x_{\mathrm{c}}, y_{\mathrm{c}})$ of the centre of the rod and its orientation $\phi$ with respect to the channel direction $x_2$ (Fig. S8a). Similar to hard sphere models in the statistical theory of solids and fluids, the hard rod is defined as a body that does not interact with the channel but cannot penetrate the walls of the channel either, i.e. infinitely strong repulsion to any overlap between the channel walls and the hard rod. The energy $E(x_{\mathrm{c}}, \phi)$ of a microstate (being independent of $y_{\mathrm{c}}$) can then be specified in terms of two cases: (i) the narrow channel with $\bar{w} \leq 1$, and (ii) the wide channel with $\bar{w} > 1$, where $\bar{w} \equiv w/L$. The energy of the microstates for the narrow channel is given for all $y_{\mathrm{c}}$ as

$$E(\bar{x}_c, \phi) \equiv \begin{cases} 0 & |\bar{x}_c| \le \bar{w}/2 \text{ and } |\phi| \le \phi_c \\ \infty & \text{otherwise,} \end{cases} \tag{4.1}$$

where $\bar{x}_c \equiv x_c/L$, with geometrical constructions specifying that the maximum angular misalignment of the rod with the channel is $\phi = \phi_c = \sin^{-1}(\bar{w} - 2\bar{x}_c)$; see Fig. S8a. Similarly, the energy of the microstates for the wide channel follows as

$$E(\bar{x}_c, \phi) \equiv \begin{cases} 0 & |\bar{x}_c| \le (\bar{w} - 1)/2 \\ 0 & (\bar{w} - 1)/2 < |\bar{x}_c| \le \bar{w}/2 \quad \text{and} \quad |\phi| \le \phi_c \\ \infty & \text{otherwise.} \end{cases} \tag{4.2}$$

Without specifying the mechanism for the fluctuations of the hard-rod over the available microstates, we shall make the ansatz that the equilibrium macrostate of the hard rod maximises the entropy of the microstates subject to appropriate constraints that will be specified subsequently. We proceed to derive this macrostate.

Since the energy $E$ is independent of $y_c$, all microstates translated along the $x_2$ −direction are equally probable. Thus, without loss of generality for the purposes of entropy maximisation, we can specify a microstate by $(\bar{x}_c, \phi)$, with $P(\bar{x}_c, \phi)$ denoting the joint probability of the hard rod having its centre at $\bar{x}_c$ and an orientation $\phi$. This joint probability can be rewritten in terms of the conditional probability $P(\phi|\bar{x}_c)$ of the rod having an orientation $\phi$ when its centre is located at $\bar{x}_c$ as

$$P(\bar{x}_c, \phi) = P(\bar{x}_c)P(\phi|\bar{x}_c), \tag{4.3}$$

where $P(\bar{x}_c) \equiv \sum_{(\phi|\bar{x}_c)} P(\bar{x}_c, \phi)$ is the probability of the centre of the rod being located at $\bar{x}_c$. The total entropy of the system is then defined as

$$I_T \equiv -\sum_{\bar{x}_c, \phi} P(\bar{x}_c, \phi) \ln P(\bar{x}_c, \phi) = I_x + \sum_{\bar{x}_c} P(\bar{x}_c) I_{(\phi|\bar{x}_c)} = I_x + I_\phi \,, \tag{4.4}$$

where $I_x \equiv -\sum_{\bar{x}_c} P(\bar{x}_c) \ln P(\bar{x}_c)$ is the positional entropy of the rod, while $I_{(\phi|\bar{x}_c)} \equiv -\sum_\phi P(\phi|\bar{x}_c) \ln P(\phi|\bar{x}_c)$ is the orientational entropy of the rod with centre located at $\bar{x}_c$, and $I_\phi$ is the total orientational entropy of the rod.

To determine the probability distributions, we need to maximise $I_T$ subject to appropriate constraints which include $\sum_{\bar{x}_c} P(\bar{x}_c) = 1$ and $\sum_\phi P(\phi|\bar{x}_c) = 1$. To understand any additional constraints, we follow the procedure first introduced by Gibbs in defining the canonical ensemble; see for example Kuzemksy [26]. Consider an assembly comprising $N_A$ number of copies of the system, each in one of the microstates that the system attains at equilibrium. This large assembly is assumed to be isolated from its surroundings. Therefore, the energy of the assembly remains constant such that $\sum_{\bar{x}_c, \phi} P(\bar{x}_c, \phi) E(\bar{x}_c, \phi) = \bar{E}$, where $\bar{E} N_A$ is the total energy of the isolated assembly. Thus, $I_T$ needs to be maximised with the above three constraints imposed by Lagrange multipliers $\lambda_i > 0$ $(i = 1, 2, 3)$ such that

$$dI_x + \sum_{\bar{x}_\mathrm{c}} P(\bar{x}_\mathrm{c}) dI_{(\phi|\bar{x}_\mathrm{c})} + \sum_{\bar{x}_\mathrm{c}} I_{(\phi|\bar{x}_\mathrm{c})}\, dP(\bar{x}_\mathrm{c}) - \lambda_1 \left[ \sum_{\bar{x}_\mathrm{c},\phi} P(\phi|\bar{x}_\mathrm{c}) E(\bar{x}_\mathrm{c},\phi) dP(\bar{x}_\mathrm{c}) + \sum_{\bar{x}_\mathrm{c},\phi} P(\bar{x}_\mathrm{c}) E(\bar{x}_\mathrm{c},\phi) dP(\phi|\bar{x}_\mathrm{c}) \right] - \lambda_2 \sum_{\bar{x}_\mathrm{c}} dP(\bar{x}_\mathrm{c}) - \lambda_3 \sum_{\phi} dP(\phi|\bar{x}_\mathrm{c}) = 0. \tag{4.5}$$

Noting that the arbitrary variations $dP(\bar{x}_\mathrm{c})$ and $dP(\phi|\bar{x}_\mathrm{c})$ are independent, the above equations split into two independent equations

$$\sum_{\bar{x}_\mathrm{c}} P(\bar{x}_\mathrm{c}) dI_{(\phi|\bar{x}_\mathrm{c})} - \lambda_1 \sum_{\bar{x}_\mathrm{c},\phi} P(\bar{x}_\mathrm{c}) E(\bar{x}_\mathrm{c},\phi) dP(\phi|\bar{x}_\mathrm{c}) - \lambda_3 \sum_{\phi} dP(\phi|\bar{x}_\mathrm{c}) = 0, \tag{4.6}$$

and

$$dI_x + \sum_{\bar{x}_\mathrm{c}} I_{(\phi|\bar{x}_\mathrm{c})}\, dP(\bar{x}_\mathrm{c}) - \lambda_1 \sum_{\bar{x}_\mathrm{c},\phi} P(\phi|\bar{x}_\mathrm{c}) E(\bar{x}_\mathrm{c},\phi) dP(\bar{x}_\mathrm{c}) - \lambda_2 \sum_{\bar{x}_\mathrm{c}} dP(\bar{x}_\mathrm{c}) = 0. \tag{4.7}$$

We shall consider both these equations in turn and first consider Eq. (4.6). Recall that $dP(\phi|\bar{x}_\mathrm{c})$ (and therefore $dI_{(\phi|\bar{x}_\mathrm{c})}$) are arbitrary and hence independent of $\bar{x}_\mathrm{c}$. It then follows that Eq. (4.6) reduces to

$$\sum_{\phi} (1 + \ln P(\phi|\bar{x}_\mathrm{c}) + \lambda_1 E(\bar{x}_\mathrm{c},\phi) + \lambda_3) dP(\phi|\bar{x}_\mathrm{c}) = 0, \tag{4.8}$$

and again, noting that $dP(\phi|\bar{x}_\mathrm{c})$ is independent of $\phi$, we have

$$1 + \ln P(\phi|\bar{x}_\mathrm{c}) + \lambda_1 E(\bar{x}_\mathrm{c},\phi) + \lambda_3 = 0\,, \qquad \forall \;\; \phi|\bar{x}_\mathrm{c}. \tag{4.9}$$

The equilibrium distribution then follows as

$$P_\mathrm{eq}(\phi|\bar{x}_\mathrm{c}) = \frac{\exp[-\lambda_1 E(\bar{x}_\mathrm{c},\phi)]}{Z_{(\phi|\bar{x}_\mathrm{c})}}, \tag{4.10}$$

where $Z_{(\phi|\bar{x}_\mathrm{c})} \equiv \exp(\lambda_3 + 1)$ is the orientational partition function of the rod located at $\bar{x}_\mathrm{c}$, and the corresponding equilibrium value of the entropy $I_{(\phi|\bar{x}_\mathrm{c})}$ is given by $S_{(\phi|\bar{x}_\mathrm{c})} \equiv -\sum_\phi P_\mathrm{eq}(\phi|\bar{x}_\mathrm{c}) \ln P_\mathrm{eq}(\phi|\bar{x}_\mathrm{c})$.

It now remains to give explicit expressions for $P_\mathrm{eq}(\phi|\bar{x}_\mathrm{c})$ by eliminating the Lagrange multiplier in Eq. (4.10). Recalling that $E(\bar{x}_\mathrm{c},\phi) = 0$ or $\infty$, the conditional probabilities

$$P_\mathrm{eq}(\phi|\bar{x}_\mathrm{c}) = \begin{cases} 0 & E(\bar{x}_\mathrm{c},\phi) = \infty \\ 1/Z_{(\phi|\bar{x}_\mathrm{c})} & E(\bar{x}_\mathrm{c},\phi) = 0\,, \end{cases} \tag{4.11}$$

with $\sum_\phi P_\mathrm{eq}(\phi|\bar{x}_\mathrm{c}) E(\bar{x}_\mathrm{c},\phi) = 0, \forall\; \bar{x}_\mathrm{c}$. Thus, $S_{(\phi|\bar{x}_\mathrm{c})} = \ln Z_{(\phi|\bar{x}_\mathrm{c})}$, and this entropy can be calculated as follows. Assuming that the rod can assume $N_\phi \gg 1$ equally spaced orientational states over $-\pi/2 \le \phi \le \pi/2$, it follows that for the narrow channel

$$P_\mathrm{eq}(\phi|\bar{x}_\mathrm{c}) = \begin{cases} \dfrac{\pi}{2\phi_\mathrm{c} N_\phi} & |\bar{x}_\mathrm{c}| \le \bar{w}/2 \\ 0 & \text{otherwise}, \end{cases} \tag{4.12}$$

while for the wide channel

$$P_{\mathrm{eq}}(\phi|\bar{x}_{\mathrm{c}}) = \begin{cases} \dfrac{1}{N_\phi} & |\bar{x}_{\mathrm{c}}| \le (\bar{w}-1)/2 \\ \dfrac{\pi}{2\phi_{\mathrm{c}} N_\phi} & (\bar{w}-1)/2 < |\bar{x}_{\mathrm{c}}| \le \bar{w}/2 \\ 0 & \text{otherwise.} \end{cases} \quad (4.13)$$

The orientational partition function of the rod located at $\bar{x}_{\mathrm{c}}$ in the narrow channel then follows as $Z_{(\phi|\bar{x}_{\mathrm{c}})} = 2\phi_{\mathrm{c}} N_\phi/\pi \;\; \forall\, \bar{x}_{\mathrm{c}}$, while in the wide channel

$$Z_{(\phi|\bar{x}_{\mathrm{c}})} = \begin{cases} N_\phi & |\bar{x}_{\mathrm{c}}| \le (\bar{w}-1)/2 \\ \dfrac{2\phi_{\mathrm{c}} N_\phi}{\pi} & (\bar{w}-1)/2 < |\bar{x}_{\mathrm{c}}| \le \bar{w}/2. \end{cases} \quad (4.14)$$

Now, returning to Eq. (4.7), we rewrite (4.7) using the equilibrium values of $P(\phi|\bar{x}_{\mathrm{c}})$ and $I_{(\phi|\bar{x}_{\mathrm{c}})}$ such that

$$-\sum_{\bar{x}_{\mathrm{c}}} \left[ 1 + \ln P(\bar{x}_{\mathrm{c}}) - S_{(\phi|\bar{x}_{\mathrm{c}})} + \lambda_1 \sum_{\bar{x}_{\mathrm{c}},\phi} P_{\mathrm{eq}}(\phi|\bar{x}_{\mathrm{c}}) E(\bar{x}_{\mathrm{c}},\phi) + \lambda_2 \right] dP(\bar{x}_{\mathrm{c}}) = 0. \quad (4.15)$$

Noting that $\sum_\phi P_{\mathrm{eq}}(\phi|\bar{x}_{\mathrm{c}}) E(\bar{x}_{\mathrm{c}},\phi) = 0$, and that the variations $dP(\bar{x}_{\mathrm{c}})$ are arbitrary and independent of $\bar{x}_{\mathrm{c}}$, the equilibrium probability distribution $P_{\mathrm{eq}}(\bar{x}_{\mathrm{c}})$ follows as

$$P_{\mathrm{eq}}(\bar{x}_{\mathrm{c}}) = \frac{\exp\left[S_{(\phi|\bar{x}_{\mathrm{c}})}\right]}{Z_{\bar{x}_{\mathrm{c}}}} = \frac{Z_{(\phi|\bar{x}_{\mathrm{c}})}}{Z_{\bar{x}_{\mathrm{c}}}}, \quad (4.16)$$

where $Z_{\bar{x}_{\mathrm{c}}} \equiv \exp(\lambda_2 + 1)$ is the positional partition function of the rod, with the equilibrium value of $I_x$ given as $S_x \equiv -\sum_{\bar{x}_{\mathrm{c}}} P_{\mathrm{eq}}(\bar{x}_{\mathrm{c}}) \ln P_{\mathrm{eq}}(\bar{x}_{\mathrm{c}}) = \ln Z_{\bar{x}_{\mathrm{c}}}$. The equilibrium probability distribution of the orientations the rod assumes then follows directly as $P_{\mathrm{eq}}(\phi) \equiv \sum_{(\bar{x}_{\mathrm{c}}|\phi)} P_{\mathrm{eq}}(\bar{x}_{\mathrm{c}},\phi)$, where $P_{\mathrm{eq}}(\bar{x}_{\mathrm{c}},\phi) \equiv P_{\mathrm{eq}}(\bar{x}_{\mathrm{c}}) P_{\mathrm{eq}}(\phi|\bar{x}_{\mathrm{c}})$.

Explicit expressions for $P_{\mathrm{eq}}(\bar{x}_{\mathrm{c}})$ can again be obtained by assuming a discretisation for the positional microstates. Here, we assume that the rod centre can assume discrete states equally spaced a distance $\Delta w$ apart, so that the centre of the rod can exist in $\bar{w}/\Delta\bar{w}$ positions along the channel width, where $\Delta\bar{w} \equiv \Delta w/L \ll 1$. For the narrow channels, $Z_{\bar{x}_{\mathrm{c}}}$ is then given by

$$Z_{\bar{x}_{\mathrm{c}}} = \frac{4N_\phi}{\pi} \sum_{k=1}^{\frac{\bar{w}}{2\Delta\bar{w}}} \sin^{-1}\left(\bar{w} - 2k\frac{\Delta\bar{w}}{\bar{w}}\right) + \frac{2N_\phi}{\pi} \sin^{-1}(\bar{w}), \quad (4.17)$$

while for the wide channels

$$Z_{\bar{x}_{\mathrm{c}}} = N_\phi \left(\frac{\bar{w}-1}{\Delta\bar{w}}\right) + \frac{4N_\phi}{\pi} \sum_{k=1}^{\frac{\bar{w}}{2\Delta\bar{w}}} \sin^{-1}\left(1 - 2k\frac{\Delta\bar{w}}{\bar{w}}\right). \quad (4.18)$$

To illustrate these predictions, we include in Fig. S8b the joint probability density functions $p_{\mathrm{eq}}(\hat{x}_{\mathrm{c}},\phi)$ for four selected normalised channel widths $\bar{w}$, where $\hat{x}_{\mathrm{c}} \equiv x_{\mathrm{c}}/w = \bar{x}_{\mathrm{c}}/\bar{w}$. This probability density is defined as

$$p_{\mathrm{eq}}(\hat{x}'_{\mathrm{c}},\phi') d\hat{x}_{\mathrm{c}} d\phi = P_{\mathrm{eq}}\left[\left(\hat{x}'_{\mathrm{c}} - \frac{d\hat{x}_{\mathrm{c}}}{2}, \phi' - \frac{d\phi}{2}\right) \le (\hat{x}_{\mathrm{c}},\phi) \le \left(\hat{x}'_{\mathrm{c}} + \frac{d\hat{x}_{\mathrm{c}}}{2}, \phi' + \frac{d\phi}{2}\right)\right], \quad (4.19)$$

where $\pi/N_\phi \ll d\phi \ll \pi$ and $\Delta\bar{w} \ll d\bar{x}_{\mathrm{c}} \ll 1$ such that

$$\int_{-w/2}^{w/2}\int_{-\pi/2}^{\pi/2} p_{\text{eq}}(\hat{x}_{\text{c}}, \phi) d\hat{x}_{\text{c}} d\phi = 1. \quad (4.20)$$

This definition ensures that $p_{\text{eq}}(\hat{x}_c, \phi)$ is independent of the choice of $\Delta\bar{w}$ and $N_\phi$. These predictions are qualitatively similar to those of the homeostatic mechanics framework for myofibroblasts (Fig. 3). In particular, this simple hard rod model predicts nearly uniform joint probability distributions for large $\bar{w}$ with the distribution becoming increasingly peaked with decreasing $\bar{w}$. In fact, similar to the homeostatic mechanics predictions (Fig. 3), the rod can assume a wider range of orientations towards the centre of the channel (i.e. $|\bar{x}_{\text{c}}| \approx 0$), with the orientation range becoming increasingly restricted for larger values of $|\bar{x}_{\text{c}}|$. Simultaneously, also similar to the homeostatic mechanics predictions, the probability of the rod centre being located near the edges of the channel (i.e. $|\bar{x}_{\text{c}}| \to 0.5$) is lower than the rod being located towards the centre as the rod can assume fewer orientations near the channel edges. Thus, while the rod aligns with the channel when it is located near the channel edges, the probability of the rod being located near the edges is relatively low. To capture succinctly how the conflict between these two competing effects is resolved, we characterize the alignment of the rod via an order parameter completely analogous to that used to define the alignment of the cells on the fibronectin stripes, i.e.

$$\Theta_{\text{R}} = \sqrt{\langle\cos 2\phi\rangle^2 + \langle\sin 2\phi\rangle^2}\,, \quad (4.21)$$

with the ensemble averages $\langle\cos 2\phi\rangle \equiv \sum_\phi P_{\text{eq}}(\phi)\cos 2\phi$ and $\langle\sin 2\phi\rangle \equiv \sum_\phi P_{\text{eq}}(\phi)\sin 2\phi$, respectively. This parameter is plotted in Fig. 6b as a function of $\bar{w}$, and shows the increasing alignment of the rod with decreasing stripe width caused by the rods near the channel edges being more aligned.

The overall order (or disorder) of the rods within the channels can be described in terms of the equilibrium positional entropy $S_x$, orientational entropy $S_\phi \equiv \sum_{\bar{x}_{\text{c}}} P_{\text{eq}}(\bar{x}_{\text{c}}) S_{(\phi|\bar{x}_{\text{c}})}$, and the total entropy $S_{\text{T}} \equiv S_x + S_\phi$. These entropies follow directly from the above expressions for the equilibrium probabilities and are omitted here for the sake of brevity. Predictions of the variation of these entropies with $\bar{w}$ are included in Fig. 6c for the discretisation of microstates specified by $N_\phi = 314$ and $1/\Delta\bar{w} = 800$. We shall discuss the significance of these results in the context of reduced models described subsequently.

##### *4.1 Reduction to special cases*

The above analysis allows both translational and rotational freedom for the rod with a microstate specified by $(\bar{x}_{\text{c}}, \phi)$. It is instructive to also study two reduced models in order to explain how endowing additional degrees of freedom to the rod, while increasing the total disorder, results in ordering of certain observables (e.g. orientation or position). We shall consider the following three cases: (i) the full model (FM) as described above where the rod has rotational and translational degrees of freedom; (ii) a translationally constrained (TC) model, and (iii) an orientationally constrained (OC) model. We shall now proceed to describe (ii) and (iii).

In the TC model, we restrict the translation of the rod with the centre of the rod fixed at $\bar{x}_{\text{c}} = 0$, and the microstates are now described in terms of a single variable $\phi$. The equilibrium distribution $P_{\text{eq}}(\phi)$ can be determined by repeating the above analysis while setting

$$P_{\text{eq}}(\bar{x}_c) = \begin{cases} 1 & \bar{x}_{\text{c}} = 0 \\ 0 & \text{otherwise.} \end{cases} \quad (4.22)$$

The entropy maximisation will now only have two constraints viz. $\sum_\phi P(\phi)E(\bar{x}_c = 0, \phi) = \bar{E}$ and $\sum_\phi P(\phi) = 1$ with $P_{eq}(\phi)$ for the wide and narrow channels given by

$$P_{eq}(\phi) = \begin{cases} \dfrac{\pi}{2\sin^{-1}(\bar{w})N_\phi} & \bar{w} < 1 \ \text{(narrow channels)}, \\ \dfrac{1}{N_\phi} & \bar{w} \geq 1 \ \text{(wide channels)}, \end{cases} \tag{4.23}$$

while the corresponding partition functions are

$$Z_\phi = \begin{cases} \dfrac{2\sin^{-1}(\bar{w})N_\phi}{\pi} & \bar{w} < 1 \ \text{(narrow channels)}, \\ N_\phi & \bar{w} \geq 1 \ \text{(wide channels)}. \end{cases} \tag{4.24}$$

The translational entropy for the TC model is $S_x^{(\mathrm{TC})} = 0$, while $S_\phi^{(\mathrm{TC})} \equiv -\sum_\phi P_{eq}(\phi) \ln P_{eq}(\phi) = \ln Z_\phi$.

The OC model eliminates the rotational degrees of freedom of the rod, with the rod constrained to always stay aligned with the channel such that $\phi = 0$. The microstates are now described in terms of only $\bar{x}_c$. Repeating the above analysis but now setting

$$P_{eq}(\phi) = \begin{cases} 1 & \phi = 0 \\ 0 & \text{otherwise}, \end{cases} \tag{4.25}$$

gives $P_{eq}(\bar{x}_c) = \Delta\bar{w}/\bar{w}$ with $Z_{\bar{x}_c} = \bar{w}/\Delta\bar{w}$. Now, the translational entropy is $S_x^{(\mathrm{OC})} \equiv -\sum_\phi P_{eq}(\bar{x}_c) \ln P_{eq}(\bar{x}_c) = \ln Z_{\bar{x}_c}$, while $S_\phi^{(\mathrm{OC})} = 0$.

While it is obvious that the orientational order parameter $\Theta_R = 1 \ \forall \ \bar{w}$ for the OC model, the full model predicts a greater orientational ordering compared to the TC model (Fig. 6b). This apparent reduction in disorder with increasing degrees of freedom can be understood by examining the variations in the entropies $S_x$, $S_\phi$ and $S_T$ with $\bar{w}$, as shown Fig. 6c for the choice $N_\phi = 314$ and $1/\Delta\bar{w} = 800$. While the full model has a lower $S_\phi$ compared to the TC model, its total entropy $S_T$ is higher, i.e. in this case the rod has traded-off orientational entropy against translational entropy so as to attain a higher total entropy. Similarly, the OC model has a higher $S_x$ compared to the full model but a lower $S_T$, i.e. in this case the full model has traded-off translational entropy against orientational entropy so as to attain a higher total entropy. We emphasise here that while the numerical values of $S_x$, $S_\phi$ and $S_T$ depend on the choices of $N_\phi$ and $1/\Delta\bar{w}$, the relative ordering of these quantities between the three different models is not dependent on these choices. For example, irrespective of the choice of $N_\phi$ and $1/\Delta\bar{w}$, we always have $S_\phi$ for the TC model to be greater than that of the full model while $S_T$ and $S_x$ for the full model always are greater than that of the TC model.

The simple entropic analysis of the hard rod within a channel performed here serves two purposes: (i) it demonstrates the mathematical process of performing entropy maximisations in a very simple setting that generates analytical expressions, and (ii) more importantly, it illustrates that the ordering of certain observables (rod orientation in this case) takes place due to the tendency of the rod to maximise its total entropy (disorder). This is the crux of the mechanism that we hypothesise here as governing the guidance of cells on fibronectin stripes. Within the context of the full homeostatic mechanics analysis, the cells maximise their disorder subject to the spatial constraints imposed by the fibronectin stripes and the homeostatic constraint. This maximisation of total disorder results in an enhanced orientational ordering of cells with respect to the case of a homogeneous substrate even when the cell size is much smaller than the width of the fibronectin stripes.

#### Supplementary Figures

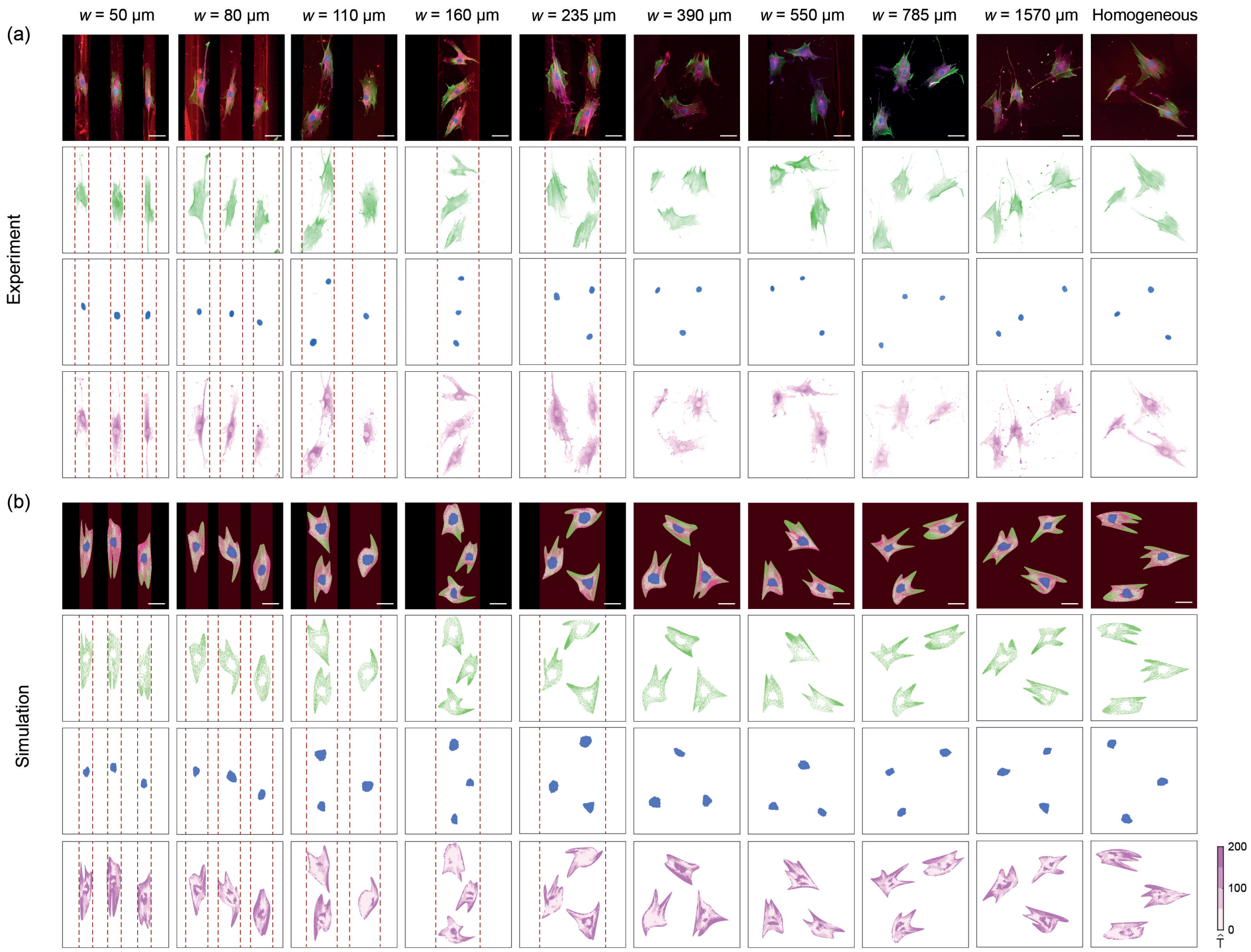

**Figure S1:** Observations and predictions across the nine fibronectin stripe widths and the control case of the homogeneous substrate. (a) Immunofluorescence images of the actin cytoskeleton (green), nucleus (blue), and vinculin (magenta) with fibronectin strained in maroon. (b) The corresponding predictions of the homeostatic mechanics framework with focal adhesions parametrised by the magnitude of the normalised traction $\widehat{\mathrm{T}}$. Scale bar = 60 μm with the width of the fibronectin stripes indicated by the dashed maroon lines for the five narrowest stripes.

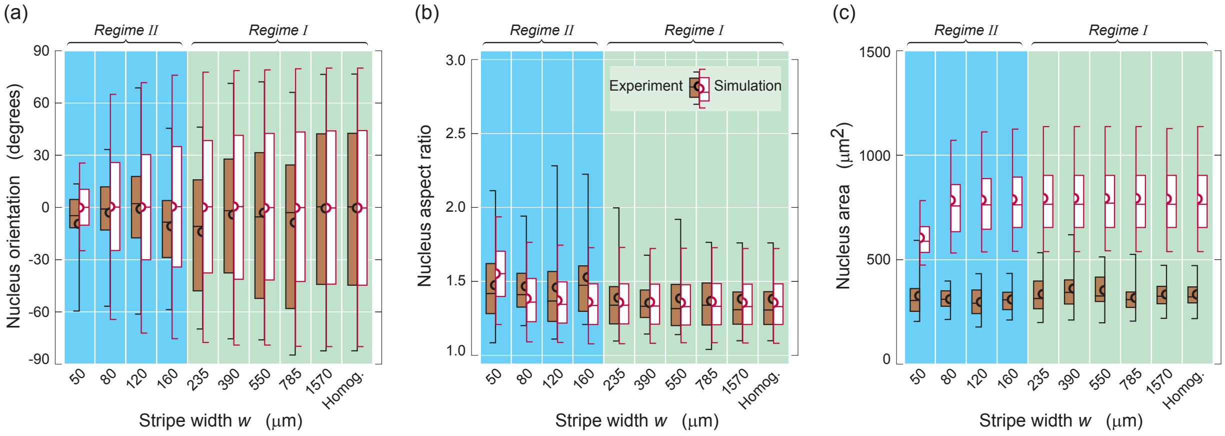

**Figure S2:** Comparison between measurements and predictions for the morphology of the nucleus. Box-and-whisker diagrams of the (a) orientation, (b) aspect ratio and (c) area of the nucleus for cells on substrates with fibronectin stripes of width $w$. The boxes show the quartiles of the distributions with the whiskers indicating the outliers in the experiments and the 5$^{\text{th}}$ and 95$^{\text{th}}$ percentiles of the distributions in the simulations. The mean of the distributions is shown by semi-circles for both the measurements and predictions. The model predicts the distribution of the nucleus orientation and aspect ratio with high fidelity, but over-predicts the area of the nucleus. We attribute the over-prediction of the nucleus area to the fact that recent studies [27] have shown the nucleus to be compressible with the volume fraction of the nucleus of cells in suspension being larger than that observed in cells on homogeneous substrates. In our calculations, we assume the nucleus to be incompressible with its volume fraction in the cell equal to that measured for myofibroblasts in their undeformed state. This results in us overestimating the nucleus volume (and therefore nucleus area) for cells on the micropatterned substrate.

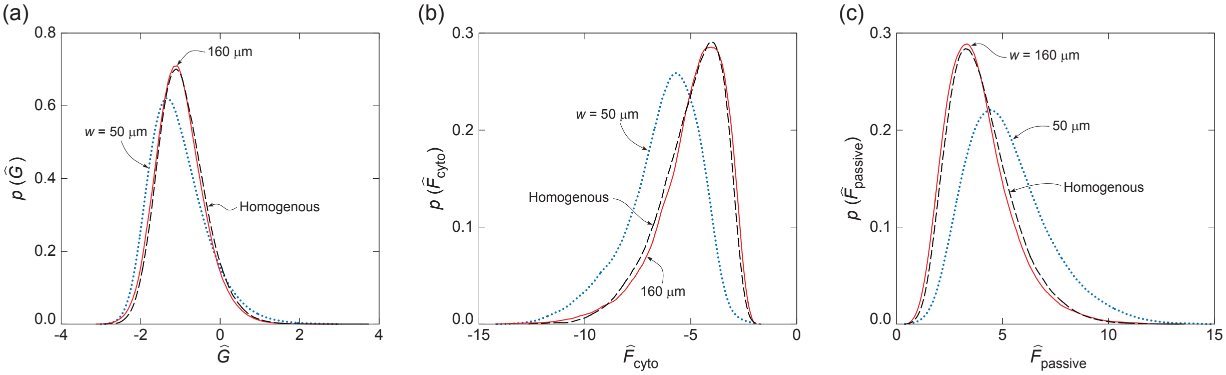

**Figure S3:** Probability density functions of cell energies. Predictions of the probability density functions of the normalised (a) Gibbs free-energy $\hat{G}$ (b) cytoskeletal free-energy $\hat{F}_{\text{cyto}}$ and (c) passive strain energy $\hat{F}_{\text{passive}}$ for myofibroblasts on substrates with fibronectin stripes of width $w$.

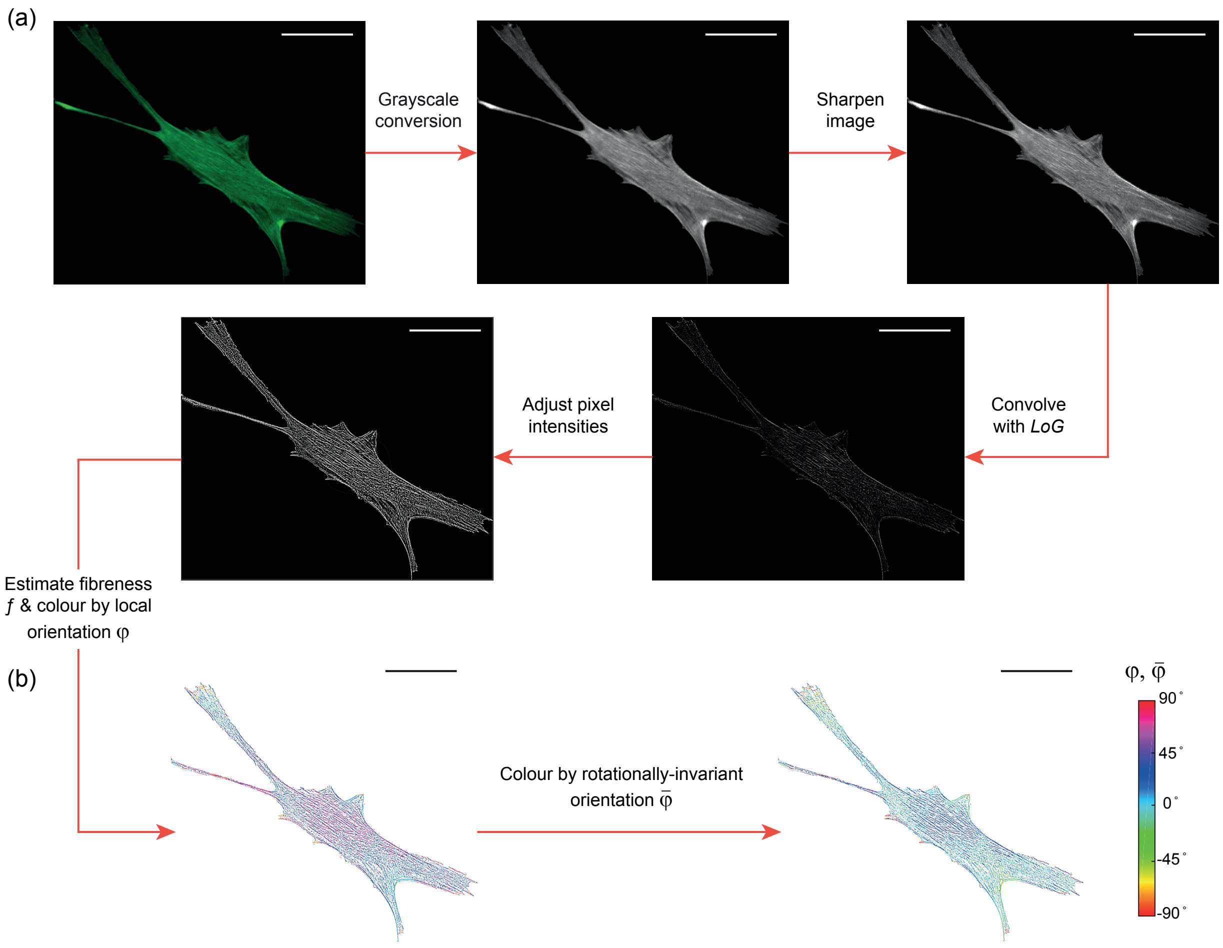

**Figure S4:** Image processing to extract spatial distributions of the orientations of stress-fibres. (a) Image processing to highlight actin fibres starting from the green channel of the immunofluorescence images, conversion to grayscale and sharpening followed by edge detection (by convolution with a Laplacian of Gaussian (*LoG*) kernel) and finally brightness adjustment to highlight the low intensity fibres. (b) Images showing the actin stress-fibres detected from "fibreness" estimation, and coloured by their respective orientations $\varphi$ with respect to the stripe direction $x_2$, and by the rotationally-invariant orientation measure $\bar{\varphi}$. The images shown here are for an imaged myofibroblast on the homogeneous substrate. Scale bar = 60 μm.

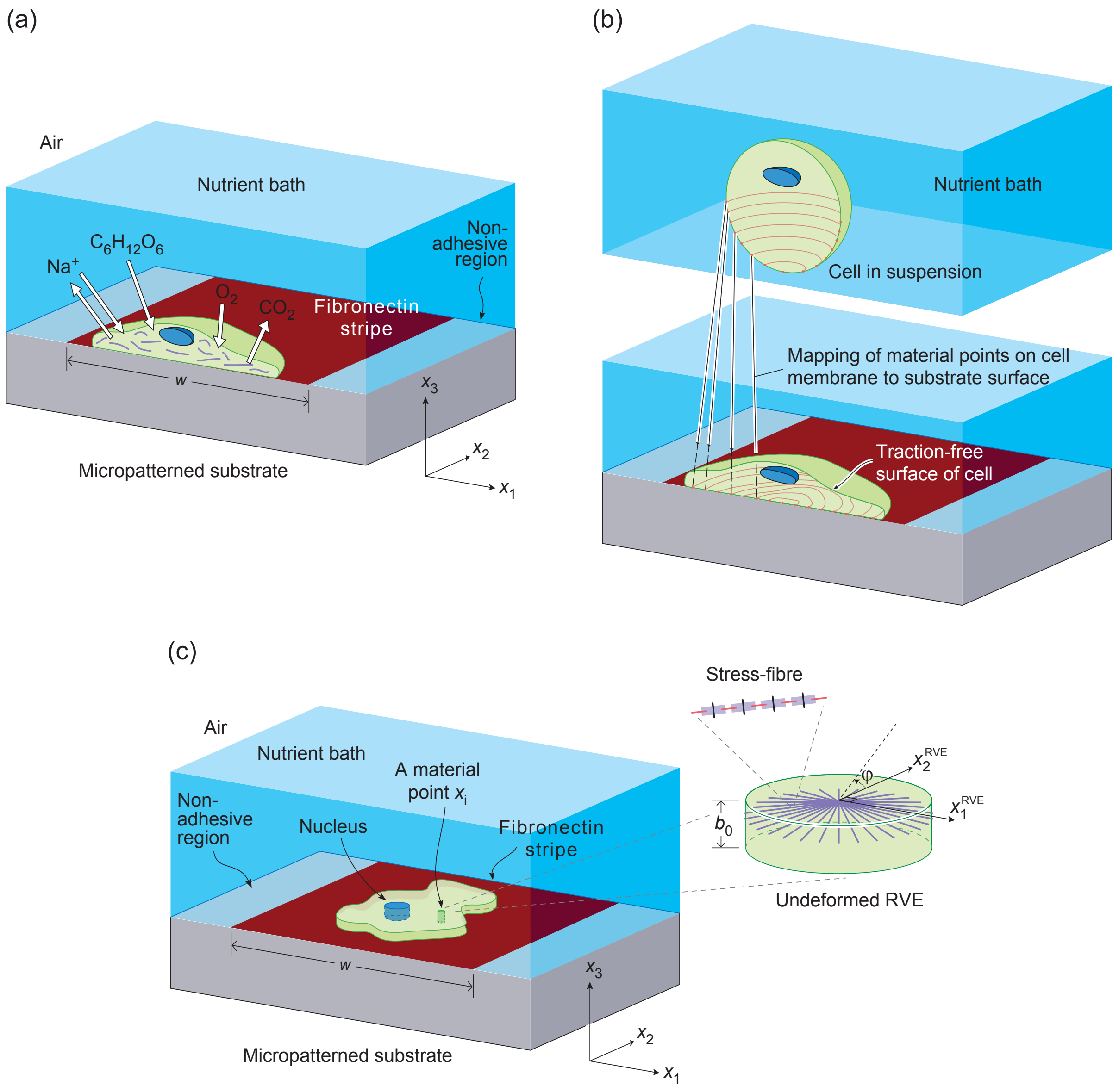

**Figure S5:** Sketches illustrating the problem of a cell on a substrate micropatterned with fibronectin stripes. (a) Sketch of a section of the experimental setup showing cells on a fibronectin stripe of width $w$ within a nutrient bath. A small selection of the species exchanged between the cells and the nutrient bath are labelled. (b) Schematic of the definition of a morphological microstate specified by the mapping of material points on the cell surface with material points on the fibronectin stripe. (c) The two-dimensional (2D) approximation of the cell on the micropatterned substrate. Sketch of the 2D morphological microstate boundary value problem comprising a 2D cell on rigid substrate and adhered with a fibronectin stripe used in the calculation of $G^{(j)}$. The co-ordinate system employed is indicated with the cell lying in the $x_1 - x_2$ plane with the through-thickness stress $\Sigma_{33} = 0$. The inset shows the cylindrical RVE along with the definition of the orientation $\varphi$ of stress-fibres.

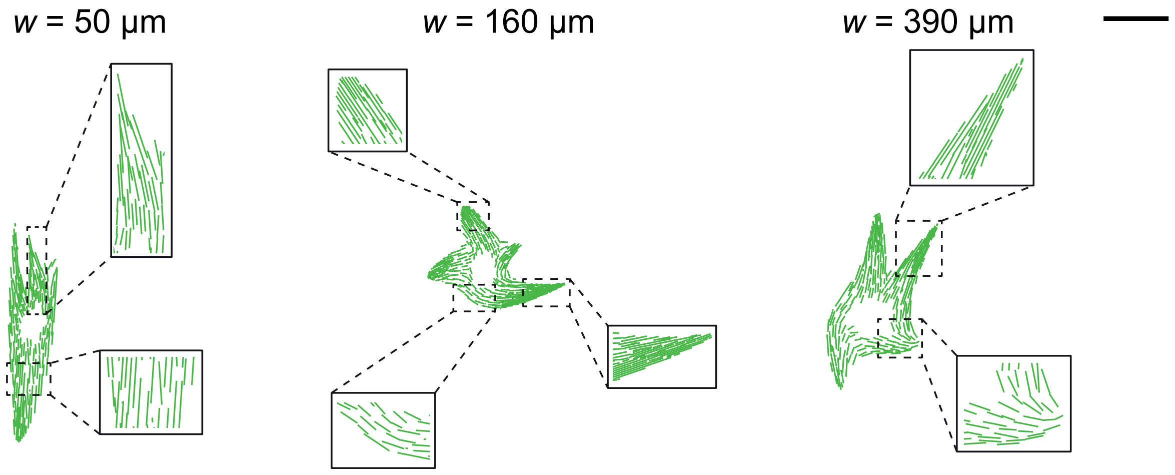

**Figure S6:** Images showing predictions of the stress-fibre distributions. Enlarged images of the actin stress-fibre structure within some of the morphological microstates shown in Fig. 1c. The insets show zoomed-in regions to give more details of the predicted stress-fibre arrangements including their orientations and spacings. Scale bar = 60 μm.

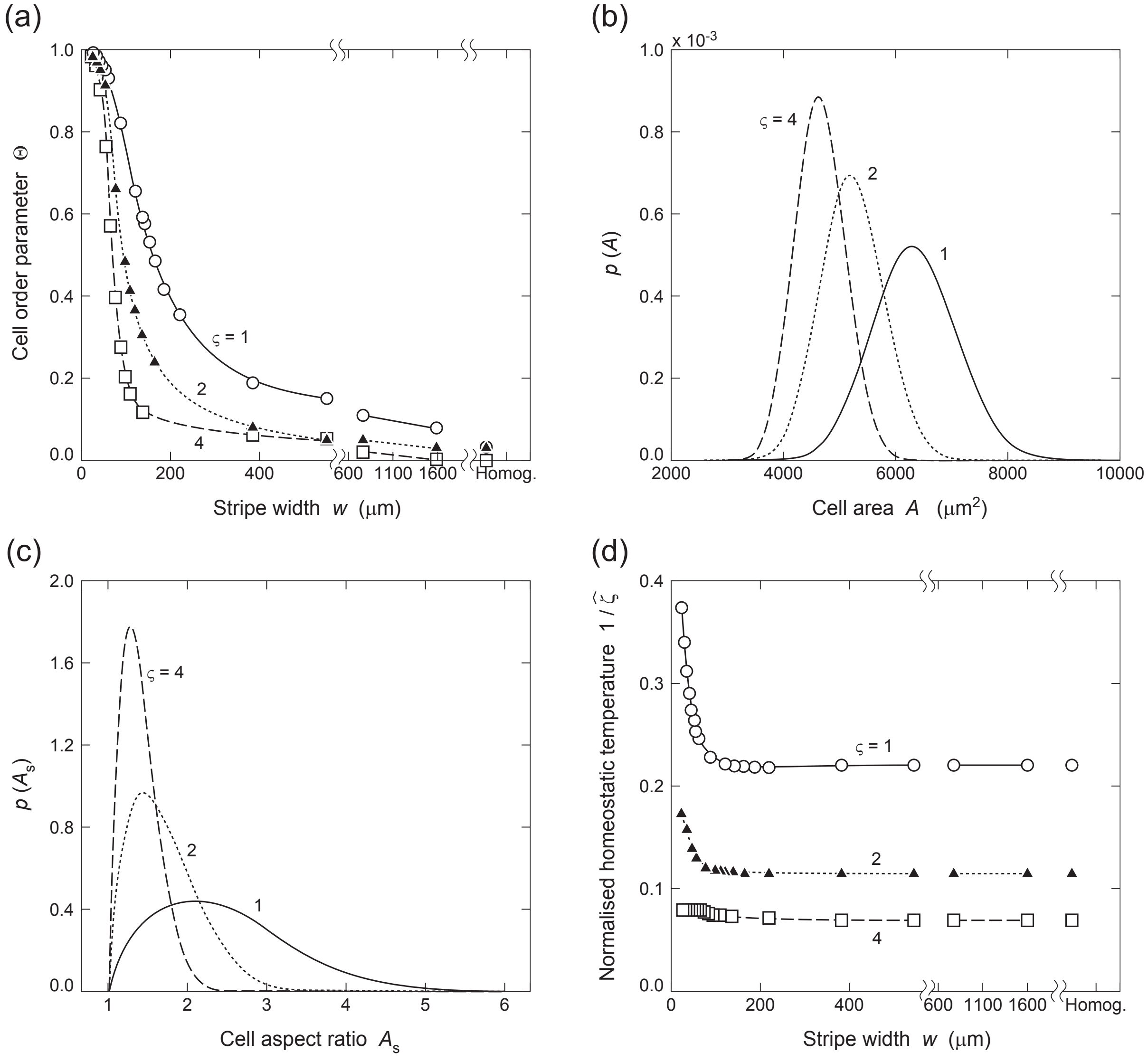

**Figure S7:** Predictions of the effect of cell stiffness on contact guidance. (a) The cell orientational order parameter Θ as a function of the fibronectin stripe width $w$. Probability density functions of the (b) cell area $A$ and (c) cell aspect ratio $A_s$. (d) Variation of the normalised homeostatic temperature $1/\hat{\zeta}$ with $w$. Results in each case are shown for three cell stiffnesses parameterised by $\varsigma$ with $\varsigma = 1$ corresponding to the reference case of myofibroblasts.

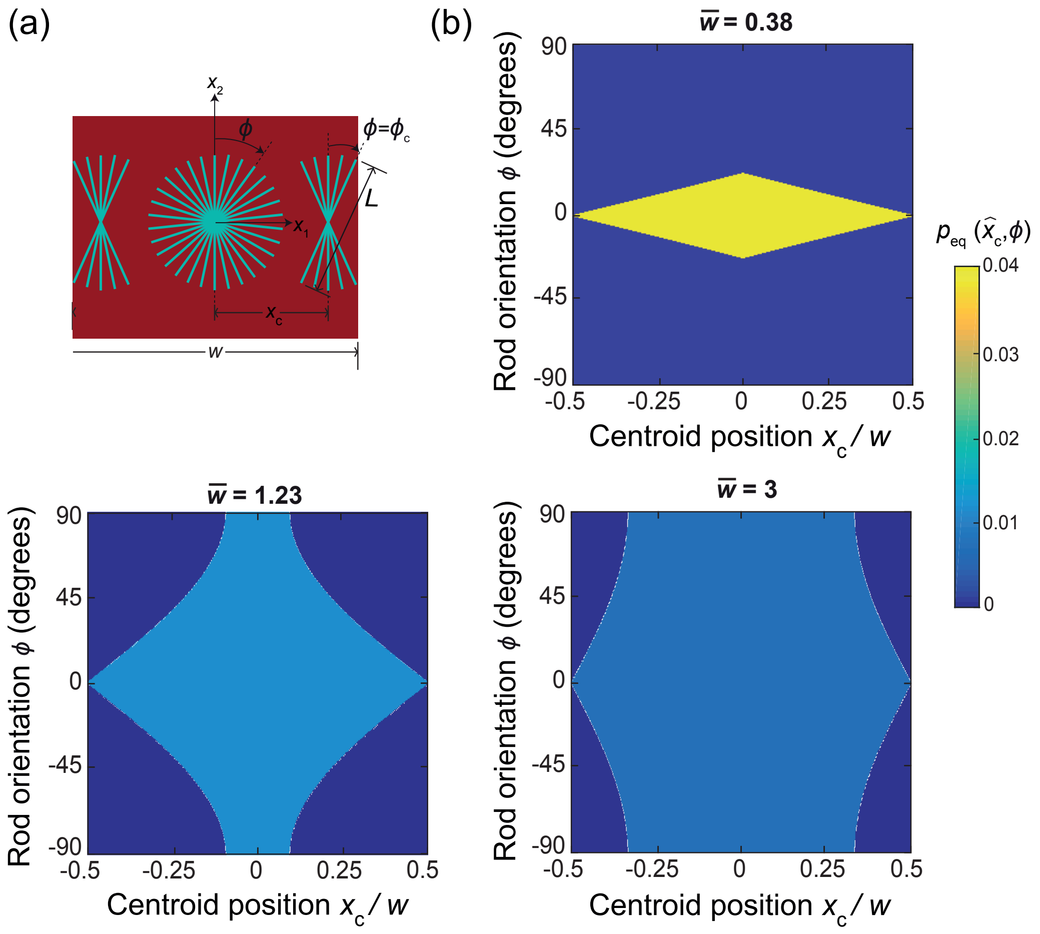

**Figure S8:** Entropic alignment of a hard rod within a channel of width $w$. (a) Sketch defining the problem along with the co-ordinate system and the co-ordinates ($x_c$, $\phi$) that define a microstate. The maximum angle $\phi = \phi_c$ that the rod can be misaligned by when near the edge of the channel is also illustrated. (b) Joint probability distributions $p_{eq}(\hat{x}_c, \phi)$ for the position and orientation of a hard rod within a channel of width $w$. Results are shown for 3 selected values of the normalised channel width $\overline{w} \equiv w/L$ with the normalised position of the rod centre defined as $\hat{x}_c \equiv x_c/w$.
